# Plasmodesmal closure elicits stress responses

**DOI:** 10.1101/2024.05.08.593115

**Authors:** Estee E. Tee, Andrew Breakspear, Diana Papp, Hannah R. Thomas, Catherine Walker, Annalisa Bellandi, Christine Faulkner

## Abstract

Plant cells are connected to their neighbors via plasmodesmata facilitating the exchange of nutrients and signaling molecules. During immune responses, plasmodesmata close, but how this contributes towards a full immune response is unknown. To investigate this, we developed two transgenic lines with which we could induce plasmodesmal closure independently of immune elicitors, using the over-active CALLOSE SYNTHASE3 allele *icals3m* and the C-terminus of PDLP1 to drive callose deposition at plasmodesmata. Induction of plasmodesmal closure increased the expression of stress responsive genes, salicylic acid accumulation and resistance to *Pseudomonas syringae* DC3000. More homogeneous plasmodesmal closure using *icals3m* also led to the accumulation of starch and sugars, decreased leaf growth, as well as hypersusceptibility to *Botrytis cinerea*. Based on the profile of responses, we conclude that plasmodesmal closure itself activates stress signaling, raising questions of what signals mediate this response and whether these responses occur in all circumstances when plasmodesmata close.

## Introduction

Plants are multicellular organisms, with almost every plant cell cytoplasmically connected to its neighbors. While most plant cells are equipped with machinery to respond autonomously to a range of stress signals, optimal responses involve the regulation of this cytoplasmic connectivity between cells. This is particularly evident in immune signaling; most cells produce the receptors that perceive microbial threats and can activate appropriate responses, but the degree of connectivity to neighboring cells is a critical part of the full immune response (Lee et al. 2011; Faulkner et al. 2013). Indeed, if plants cannot regulate the cytoplasmic connectivity between cells, they are more susceptible to infection by a range of pathogenic microbes. Despite this observation, we don’t yet understand how cell-to-cell connectivity contributes to an overall immune response.

Cytoplasmic connectivity between plant cells is established via membrane-lined bridges called plasmodesmata. Plasmodesmata are dynamic structures and the oscillation in their aperture, i.e., closing and opening, allows control over the flux of soluble molecules between cells. We broadly assume that many molecules that move through plasmodesmata, including plant hormones, mRNAs, and proteins, are information carriers and that aperture changes thus impact cell-to-cell communication. Indeed, if an immune response involves the production of soluble defense-associated molecules, their passage through plasmodesmata could transmit critical information to neighboring (or distant) naïve cells.

Plant immune responses include a decrease in plasmodesmal aperture, which reduces cell-to-cell movement of molecules (as recently reviewed by Wang et al. 2021b; German et al. 2023; Alazem and Burch-Smith 2024). Plasmodesmal aperture is regulated via callose deposition and degradation at the plasmodesmal neck by callose synthases (CalS) and β-glucanases respectively (Levy et al. 2007; Vatén et al. 2011). The decrease in plasmodesmal aperture in response to stress is a process activated by signaling cascades mediated by specific machinery. For example, LYSM-CONTAINING GPI-ANCHORED PROTEIN 2 (LYM2; Faulkner et al. 2013) and CALMODULIN-LIKE 41 (CML41; Xu et al. 2017) specifically mediate plasmodesmal responses to the microbial elicitors chitin and flg22 respectively, and PLASMODESMATA LOCATED PROTEIN 5 (PDLP5) integrates these responses, as well as plasmodesmal responses triggered by the defense hormone salicylic acid (SA; Wang et al. 2013; Tee et al. 2023b). The observation that *lym2* mutants cannot close their plasmodesmata in response to chitin but execute normal chitin-triggered mitogen-activated protein kinase (MAPK) activation and apoplastic ROS production (Faulkner et al. 2013) indicates that plasmodesmal signaling cascades can act independently of other immune responses. This independence of response suggests that whether plasmodesmata are open or closed might be informed by another layer of cellular regulation.

It has been observed that plasmodesmal permeability decreases upon PAMP perception within 30 min (Xu et al. 2017) and is still detected after 24 hours (Lim et al. 2016; Li et al. 2021). Both PAMP perception and pathogen infection initiate a wide array of defense responses making it challenging to identify the specific contribution of plasmodesmal closure to immunity. Further, in the context of an infection, many pathogens deploy effectors targeting plasmodesmata that suppress plasmodesmal closure (e.g. Aung et al, 2020; Tomczynska et al. 2020; Li et al. 2021; Ohtsu et al. 2024), further complicating the analysis of the role of plasmodesmata in host immune execution in an infection context.

Whether plasmodesmata can close or not during immune responses determines whether a full defense response can be executed but untangling how plasmodesmal closure contributes to overall immunity is a complex problem. To begin to address this we have simplified the question and asked what cellular responses are triggered by plasmodesmal closure, and how closing plasmodesmata affects elements of an immune response. We generated two genetic tools with which we can induce plasmodesmal closure independently of an immune or stress elicitor. By identifying what responses are trigged by plasmodesmal closure in both tools, we found that plasmodesmal closure itself instigates a subset of stress responses including transcriptional reprogramming and salicylic acid (SA) production. This raises further questions regarding how this contributes to the orchestration of immune responses, how plasmodesmal closure triggers stress responses and whether these stress responses occur during the broad range of physiological processes during which plasmodesmata close.

## Results

### Genetic tools induce plasmodesmal closure independent of a physiological elicitor

To manipulate plasmodesmal closure independently of external signals we generated two transgenic lines in which estradiol application can induce callose deposition at plasmodesmata. First, we exploited the *icals3m* overactive *CalS3* allele (Vatén et al. 2011) that enhances callose deposition at plasmodesmata; *icals3m* has been used extensively as a tool to understand symplastic connection in development (e.g., Paterlini et al. 2021; Ross-Elliott et al. 2017; Sevilem et al. 2012; Wu et al. 2016; Yadav et al. 2014) but less is known about its effect on immunity. As a callose synthase, this enzyme acts terminally in the plasmodesmal-associated callose deposition pathway. Secondly, we utilized the observation that a synthetic construct of the fluorescent protein mCherry fused between the PDLP1 signal peptide, and the PDLP1 transmembrane domain and cytoplasmic tail, also promotes callose deposition at plasmodesmata (Caillaud et al. 2014). We fused these two transgenes independently to the *XVE* chimeric transcription activator and *LexA* operator (Zuo et al. 2000) and transformed them into *Arabidopsis thaliana* Col-0 to create two different inducible tools, named LexA::icals3m and LexA::PD-Plug respectively. To identify functional transgenic lines, we performed qPCR, copy number analysis and/or live imaging (Supplemental Table 1; Supplemental Fig. S1).

To functionally characterize the selected LexA::PD-Plug and LexA::icals3m lines, we first measured callose deposition by quantitative live cell imaging of aniline blue stained plasmodesmal callose 24 hours (h) post estradiol treatment. At this timepoint, both LexA::PD-Plug and LexA::icals3m showed greater accumulation of callose at PD when compared to the DMSO treatment, although there was a greater response in LexA::icals3m (Fig. 1a; Supplemental Fig. S2) suggesting that this line can deposit more callose at plasmodesmata than LexA::PD-Plug. To determine whether increased callose deposition perturbed plasmodesmal permeability, we used microprojectile bombardment assays of *35S::GFP* to measure GFP flux between cells in DMSO or estradiol treated leaves. While estradiol treatment did not impact GFP flux in comparison to DMSO in wild-type Col-0 plants, we found that estradiol induction of either transgene reduced GFP flux between cells, indicating plasmodesmal aperture was reduced and cell-to-cell connectivity decreased (Fig. 1b-c; Supplemental Data Set 1). The extent of GFP flux was significantly less in induced LexA::icals3m compared to induced LexA::PD-Plug (Fig. 1c). From these results, we conclude that estradiol induces plasmodesmal callose deposition, and plasmodesmal closure in both LexA::PD-Plug and LexA::icals3m, but not Col-0.

**Figure 1.**
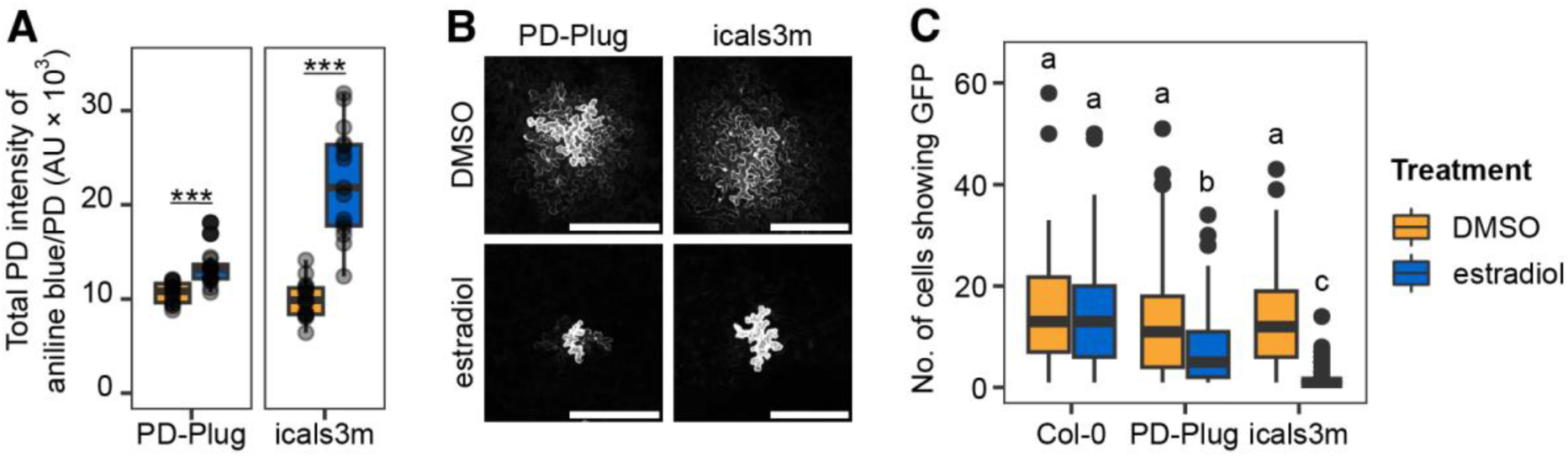
Characterization of plasmodesmal status in induced LexA::PD-Plug and LexA::icals3m tissue. **A)** Quantification of aniline blue-stained plasmodesmata-associated callose in 5-week-old plants of LexA::PD-Plug (PD-Plug) or LexA::icals3m (icals3m) 24 h post DMSO or estradiol treatment. Datapoints represent the average of the total PD intensity/plasmodesmata (PD) per image, with n ≥ 14 per genotype/treatment. Bootstrap analysis indicates significant differences between DMSO and estradiol treatment in PD-Plug and icals3m, as indicated by *** *p* < 0.001. **B)** Example micrographs of bombardment sites showing movement of GFP from transformed cells indicate cell-to-cell connectivity of uninduced (DMSO) versus restricted cell connectivity in induced (estradiol) leaves of PD-Plug and icals3m. **C)** Quantification of microprojectile bombardment data showing GFP movement into neighboring cells. Leaves of 5-week-old plants were bombarded after 24 h treatment of DMSO or estradiol; n ≥ 62 bombardment sites. Bootstrap analysis with significant differences indicated by a, b and c, with *p* < 0.01. Scale bar = 200 µM.

We noted that the data obtained from GFP mobility assays indicates that induced LexA::PD-Plug displays heterogeneous plasmodesmal closure; while there is an increase in the population of cells that exhibit plasmodesmal closure in induced tissues, a subset of the population of cells remain connected to their neighbors (Fig. 1c; Supplemental Fig. S3). This is reminiscent of the data obtained from GFP mobility assays of leaves treated with elicitors such as chitin and flg22 (Cheval et al. 2020; Tee et al. 2023). By comparison, induced LexA::icals3m displays a more homogenous response in which low variance in cell-to-cell movement is observed (Fig. 1c; Supplemental Fig. S3), similar to transgenic lines such as the PDLP1-overexpressor (Tee et al. 2023).

To further characterize the selected LexA::PD-Plug and LexA::icals3m lines, we profiled the timing of plasmodesmal callose deposition across an extended timeline. Significant plasmodesmal callose accumulation following estradiol treatment was first detected at 24 h post treatment in both LexA::PD-Plug and LexA::icals3m, and was sustained up to 72 h post treatment (Supplemental Fig. S4). Despite differences in the degree of plasmodesmal closure, these transgenic lines allow us to manipulate plasmodesmata independently of both endogenous and non-endogenous signals and investigate plant responses to plasmodesmal closure.

### Plasmodesmal closure triggers stress transcriptional responses and salicylic acid production

As pathogens are capable of manipulating plasmodesmal aperture during infection, it is difficult to determine whether plants sustain immune-associated plasmodesmal closure when challenged with a pathogen over time. Thus, we examined callose deposition in response to infection by *Pseudomonas syringae* DC3000 (*Pst* DC3000) mutant strain *hrcC^-^* that does not secrete effectors that manipulate plasmodesmal aperture. We found that plasmodesmal callose was deposited throughout the leaf at 24 h, 48 h and 72 h post infection (hpi), and significantly increased at 48 and 72 hpi (Supplemental Fig. S5). This indicates that sustained plasmodesmal closure occurs throughout an infection context.

To investigate the impact of sustained plasmodesmal closure on leaf tissue, we next characterized the transcriptional profile of this process using RNAseq, identifying transcriptional changes at 12 h, 24 h, 48 h and 72 h following DMSO or estradiol treatment in LexA::icals3m and LexA::Plug. Preliminary analysis of the data first revealed that a critical driving factor of variance was time (Supplemental Fig. S6). Therefore, all subsequent analyses were performed within single time points. We assessed the effect of estradiol relative to DMSO treatment within a genotype, identifying differentially expressed genes (DEGs). The greatest response was in Col-0 at 12 h, where estradiol led to the downregulation of over a thousand genes (Supplemental Fig. S7). We checked whether estradiol influenced plasmodesmal associated callose deposition in Col-0, but no change was detected (Supplemental Fig. S8). The presence of shared estradiol-responsive DEGs in Col-0 and the transgenic genotypes supports the possibility of an early estradiol specific response not correlated to transgene induction. The number of these shared estradiol-induced DEGs detected was dramatically reduced after 24 h (Supplemental Data Set 2; Supplemental Fig. S7), suggesting secondary transcriptional effects of estradiol are negligible after this point.

To further validate our tools, we assessed the transgene induction across the sampling period; both PD-Plug transcripts (as indicated by read counts of mCherry) and icals3m transcripts (as indicated by read counts of CalS3) were upregulated compared to controls by 12 h post estradiol treatment and remained significantly elevated at 72 h (Supplemental Fig. S9). Specifically, the PD-Plug transgene was induced to a level comparable to other highly expressed genes such as photosynthetic gene *RBCS1A* (AT1G67090; Supplemental Data Set 3). To determine that the upregulation of the transgenes did not induce ER stress from high levels of ectopic protein production, we examined the expression of genes known to be ER stress markers, defined as either ER stress sensors or unfolded protein response (UPR) effectors (Beaugelin et al. 2020). Of the 13 genes examined, only *BIP3* (AT1G09080) was significantly upregulated in estradiol treated LexA::PD-Plug and LexA::icals3m at 72 h (Supplemental Fig. S10), suggesting that transgene expression in these lines does not induce ER-stress. We also investigated gene expression changes specific to the induction of each transgene by using a likelihood-ratio test (Supplementary Data Set 4; Supplemental Fig. S11). At both 12 h and 24 h, we found no genes uniquely upregulated in the estradiol treated samples from either LexA::PD-Plug or LexA::icals3m. At 48 h, 28 and five genes were uniquely upregulated in estradiol treated LexA::PD-Plug and estradiol treated LexA::icals3m respectively. At 72h, 33 genes were differentially upregulated in estradiol treated LexA::icals3m alone; there were no genes differentially upregulated exclusively in estradiol treated LexA::PD-Plug at 72 h.

While both the *icals3m* and *PD-Plug* transgenes have different modes of action as proteins, we reasoned that the genes that are commonly up-or down-regulated following induction are associated with their common effect on plasmodesmata and likely to indicate core responses to plasmodesmal closure. To identify genes uniquely upregulated when plasmodesmata are closed, we defined two different groups for comparison: plasmodesmal state closed (estradiol treated LexA::PD-Plug and LexA::icals3m) and plasmodesmal state open (DMSO treated LexA::PD-Plug and LexA::icals3m, and all Col-0 samples). A likelihood-ratio test was performed on these two groupings at each time point to identify genes that were significantly differentially expressed only in the closed plasmodesmal state (Fig. 2a; Supplemental Fig. S12; Supplemental Data Set 5-8) and observed that the number of upregulated DEGs increased over time, suggesting that the longer plasmodesmata remain closed, the greater the cellular response. Notably, the number of differentially expressed genes that was shared between LexA::icals3m and LexA::PD-Plug in the grouping of plasmodesmal state closed was greater than the number of up-regulated DEGs specific to each line (Fig. 2a). A GO term analysis identified that genes upregulated when plasmodesmata are closed at 48 and 72 h are enriched in defensive processes (Fig. 2b; Supplemental Fig. S13). These include response to bacterium, oomycetes, chitin and fungus, regulation of systemic acquired resistance, and response to salicylic acid (SA).

**Figure 2.**
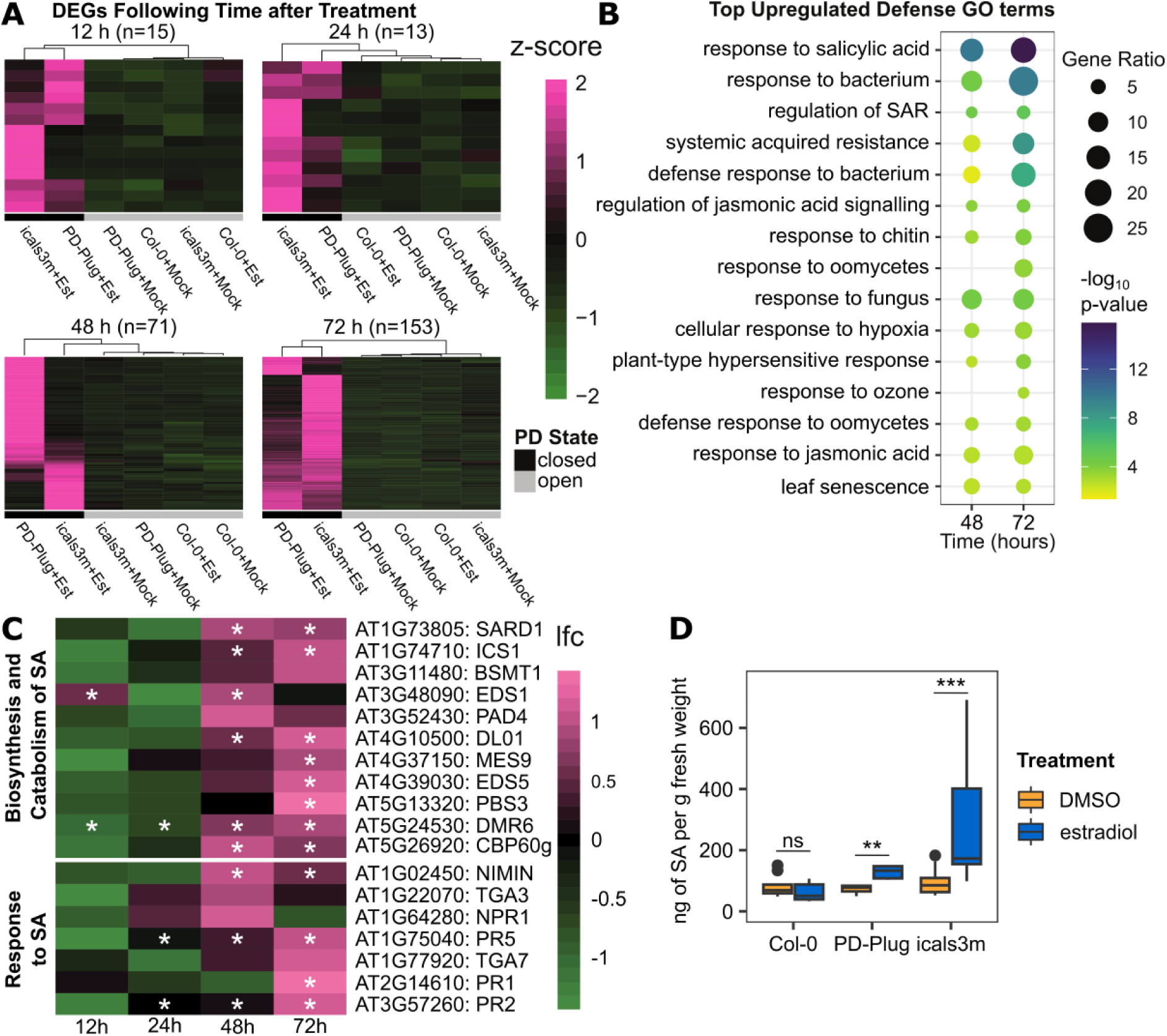
Plasmodesmal closure triggers stress transcriptional responses and salicylic acid related changes. **A)** Likelihood ratio test analysis determined number (indicated by n) of differentially expressed genes (DEGs) significantly differed between plasmodesmal status (i.e., PD State closed or open), and matches the cluster pattern of upregulated in closed state in comparison to open state at 12, 24, 48 and 72 h post treatment. Hierarchical clustering grouped together plasmodesmal state of open (i.e., estradiol treated LexA::PD-Plug and LexA::icals3m) and closed (DMSO treated LexA::PD-Plug, LexA::icals3m and Col-0, and estradiol treated Col-0). Expression normalized by row (z-score). **B)** Top 15 defense GO terms enriched over time when plasmodesmata are closed. **C)** Expression of genes related to the salicylic acid biosynthesis/catabolism and response over time when plasmodesmata are closed (i.e., in estradiol treated LexA::PD-Plug and LexA::icals3m). **D)** Quantification of salicylic acid in 5-week-old plants after 72 h DMSO or estradiol treatment, n ≥ 6 per genotype/treatment. Significant differences between treatment within a genotype analyzed using Mann-Whitney test denoted by ** p < 0.01, or *** p < 0.001.

As response to SA at 72 h was the most significantly enriched GO term in the entire analysis, we further explored this by analyzing DEGs related to SA to determine what aspects of SA signaling were impacted when plasmodesmata were closed (Fig. 2c; see Supplemental Data Set 9 for associated references; Vlot et al. 2008; Seyfferth et al. 2014; Yang et al. 2015; van Butselaar and Van den Ackerveken 2020) and found both SA biosynthesis/catabolism and SA response pathways were significantly upregulated at 48 and 72 h. To confirm that the transcriptional signature of SA is a consequence of the activation of SA synthesis when plasmodesmata are closed, we measured SA content in Col-0, LexA::PD-Plug and LexA::icalsm3 72 h post DMSO and estradiol treatment. We found induced plasmodesmal closure increased SA content in both LexA::PD-Plug and LexA::icals3m, with induced LexA::icals3m having a greatly increased SA content, being three-fold higher when compared to DMSO treatment (Fig. 2d). Previous studies have shown pathogen-induced SA accumulation is associated with the suppression of jasmonic acid (JA) signaling (e.g. Spoel et al. 2003). While GO terms associated with JA signaling and response were enriched amongst DEGs associated with 48 and 72 h PD closure (Fig. 2b, Supplemental Fig. S14), we could not quantify any differences in JA content in tissue when PD were closed (Supplemental Fig. S14). Overall, plasmodesmal closure leads to the upregulation of transcriptional defense responses, and SA synthesis and accumulation, independent of an immune stimulus.

### Plasmodesmal closure differentially impacts pathogen resistance and immune responses

Plants that lack plasmodesmata-specific immune signaling machinery and cannot close their plasmodesmata in response to pathogen infection, show increased susceptibility to different pathogenic microbes (Lee et al. 2011; Faulkner et al. 2013; Xu et al. 2017). To determine whether an inverse correlation exists for induced plasmodesmal closure we examined the susceptibility of induced, transgenic lines to the bacterial, biotrophic pathogen *Pst* DC3000. We inoculated expanded leaves of Col-0, LexA::PD-Plug and LexA::icals3m plants 72 h post DMSO or estradiol treatment (Fig. 3a) and found that induction of both transgenes reduced bacterial growth 3 days post infection (dpi). Thus, plasmodesmal closure is correlated with enhanced resistance to *Pst* DC3000. Next, we assayed for resistance to the fungal necrotroph, *Botrytis cinerea*. In contrast with *Pst* DC3000 infection, we found induction of the PD-Plug transgene had no impact on *B. cinerea* infection when compared to the disease caused in Col-0 plants, but induction of *icals3m* increased susceptibility to *B. cinerea* (Fig. 3b; Supplemental Fig. S15). This indicates that plasmodesmal closure specifically does not enhance resistance in all infection contexts.

**Figure 3.**
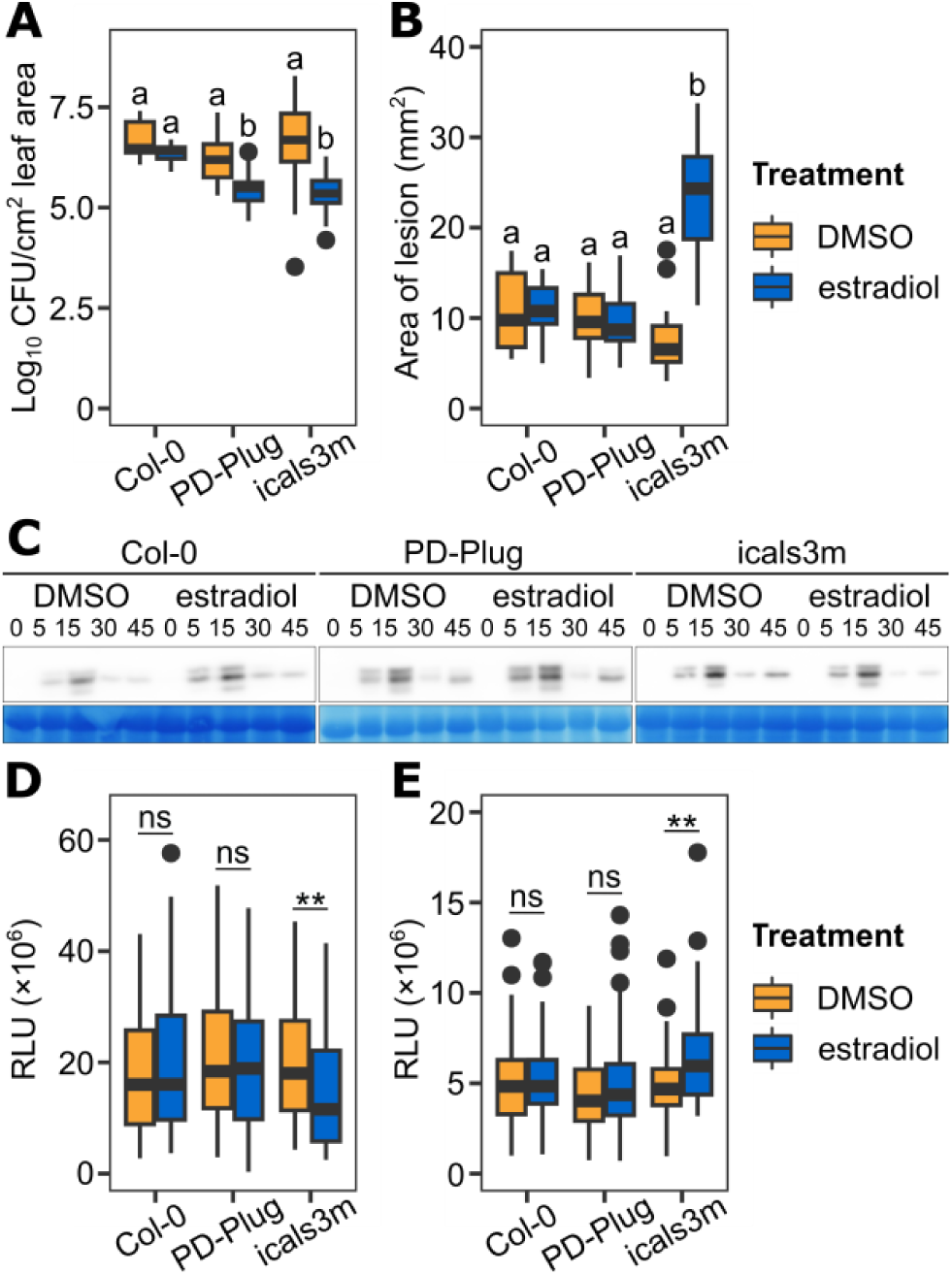
Plasmodesmal closure differentially impacts pathogen resistance and immune responses. **A)** Bacterial growth (log_10_ CFU/cm^2^ leaf area) of *Pst* DC3000 3 dpi on uninduced and induced tissue of 4–5-week-old plants of Col-0, LexA::PD-Plug (PD-Plug) and LexA::icals3m (icals3m). Tissue was treated with DMSO (uninduced) or estradiol (induced) 72 h prior to infection. 4 independent experiments were conducted, with 5 plants per treatment/genotype; N = 20. Independent factors genotype and treatment were significant, with a significant interaction between the two (ANOVA; F = 16.6, df = 2, P < 0.0001; F = 67.0, df = 1, P < 0.0001; F = 8.2, df = 2, P < 0.001). Significant differences denoted by a and b, p < 0.0001. **B)** The average area of *Botrytis cinerea* disease lesions 2 dpi in leaves of 5-week-old plants of induced and uninduced Col-0, LexA::PD-Plug and LexA::icals3m. Tissue was treated with DMSO (uninduced) or estradiol (induced) 72 h prior to infection, with N = 16-17 for all genotypes/treatment combined from three independent repeats. Independent factors genotype and treatment were significant with a significant interaction between the two (ANOVA; F = 24.4, df = 2, P < 0.0001; F = 52.7, df = 1, P < 0.0001; F = 57.0, df = 2, P < 0.0001). Significant differences denoted by a and b, p < 0.0001. **C)** Western blot detection of MAPK-activation by flg22 treatment in leaf discs from expanded leaves of 4–5-week-old plants of Col-0, LexA::PD-Plug and LexA::icals3m collected 72 h post DMSO or estradiol treatment. Minute timings of tissue harvest are denoted, and Coomassie blue staining shown below as loading controls. **D)** Average relative light units (RLU) generated from ROS production in response to flg22 in leaf discs of 5-week-old plants. Leaves were treated with estradiol or DMSO 72 h prior to flg22 treatment and data is combined from two independent experiment with N = 95 per genotype/treatment. Significant differences between treatment determined by bootstrap analysis, with significant differences denoted by **, p < 0.01. **E)** Average RLU generated from ROS production in response to flg22 in seedlings. Plants were treated with estradiol or DMSO 72 h prior to flg22 treatment, with N ≥ 80 per genotype/treatment. Significant differences between treatment determined by bootstrap analysis, with significant differences denoted by **, p < 0.01.

Overall susceptibility is driven by multiple components of host defense responses, and plasmodesmal closure is a key immune response to the pathogen associated molecular patterns (PAMPs) flg22 and chitin. To determine if plasmodesmal closure in our transgenic lines influences other PAMP responses, we assessed the activation of MAPK signaling and ROS production by flg22 in induced LexA::icals3m and LexA::Plug. When we assayed for flg22-triggered MAPK phosphorylation in mature leaves and seedlings of both lines following estradiol treatment, we saw no difference in the timing or magnitude of the response when compared to Col-0 (Fig. 3c; Supplemental Fig. S16). By contrast, while induction of *LexA::PD-Plug* did not perturb the flg22-triggered ROS burst in either mature leaves or seedlings (Fig. 3d,e), when we assayed for the same response in LexA::icals3m, induction of the transgene reduced the flg22-triggered ROS burst in mature leaves (Fig. 3d) and increased the response in seedlings (Fig. 3e). To determine whether there were differences in general basal cellular ROS levels that might explain this observation, we stained estradiol treated Col-0 and LexA::icals3m leaves with H_2_DCFDA but observed no quantitative differences between the genotypes or treatments (Supplemental Fig. S17), indicating plasmodesmal closure itself does not alter basal ROS cellular levels.

These results indicate that plasmodesmal closure differentially influences the outcomes of host interactions with different pathogenic microbes, possibly correlated with differences in pathogen lifestyle. Increased resistance to *Pst* DC3000 is not associated with enhanced MAPK activation or, given that our pathoassays were performed on mature leaves, changes in ROS burst responses. Further, our ROS data raises the possibility that plasmodesmal closure differentially interacts with immune responses in different developmental stages.

### Plasmodesmal closure induces sugar accumulation and growth defects

As plasmodesmata connect the phloem to cells in young sink tissues and allow sugar translocation, we expect that plasmodesmal closure might reduce the flux of sugars to growing tissues and limit growth. Alternatively, activation of defense processes can negatively regulate growth and therefore it is possible that plasmodesmal closure impairs growth by one or both pathways. Indeed, several transgenic lines known for constitutive plasmodesmal closure such as *PDLP1, PDLP5* and *PDLP6* overexpressors (Thomas et al. 2008; Lee et al. 2011; Li et al. 2024) exhibit a significant retardation in growth. Thus, we explored whether sustained plasmodesmal closure reduced growth in induced LexA::PD-Plug and LexA::icals3m. In 9-day old seedlings, we applied either DMSO or estradiol treatment and measured the first true leaves (Fig. 4a) six days post treatment. We found that while estradiol reduced leaf growth in both Col-0 and LexA::PD-Plug in comparison to the DMSO treated seedlings, the estradiol-induced growth reduction was greater in LexA::icals3m relative to LexA::PD-Plug and Col-0. Alongside reduced leaf size, we observed yellowing and chlorosis on the cotyledons of estradiol treated LexA::icals3m (Supplemental Fig. S18).

**Figure 4.**
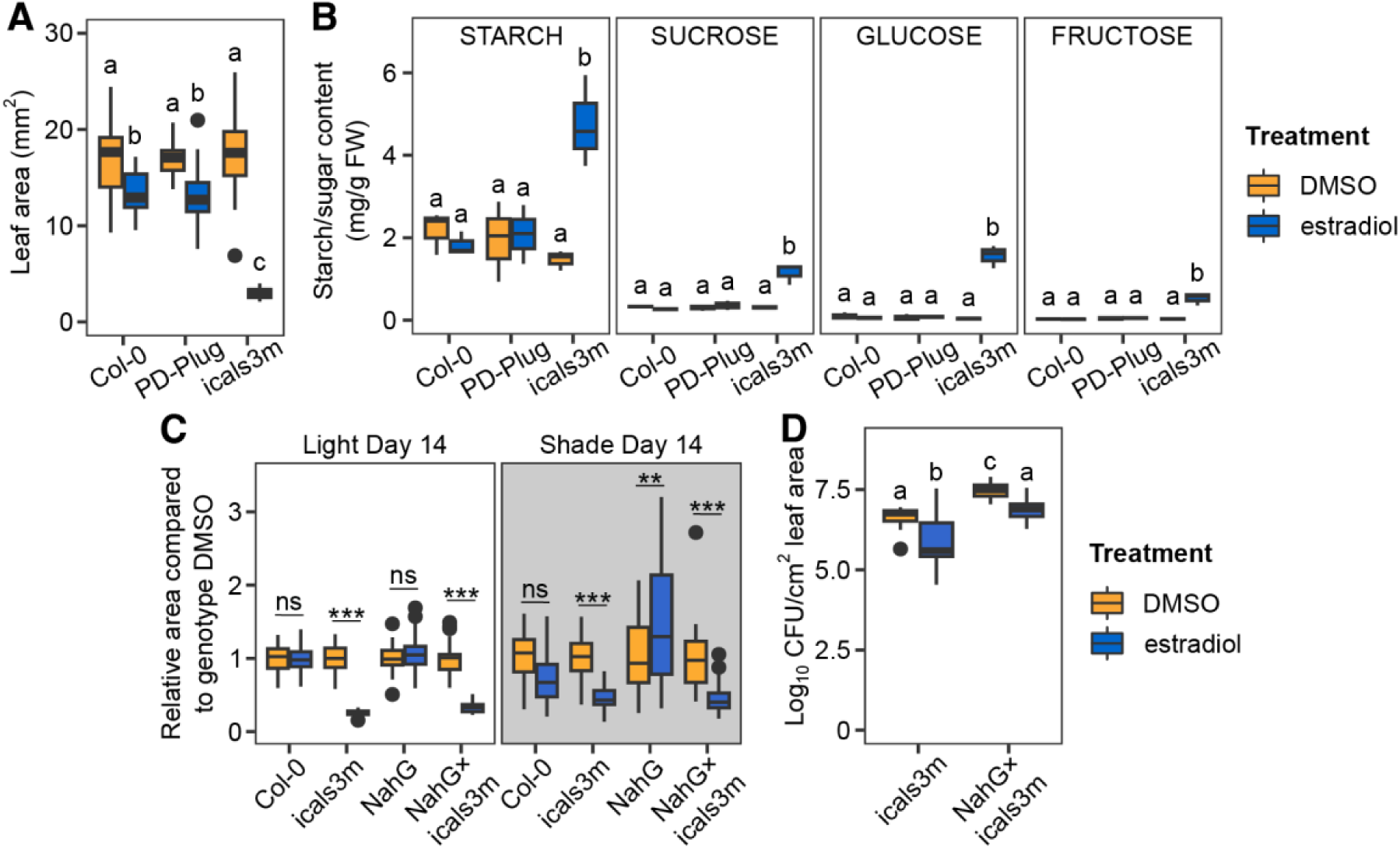
Plasmodesmal closure induces sugar accumulation and impairs leaf growth. **A)** Leaf area of first two true leaves 6 days post treatment (DMSO or estradiol) of Col-0, LexA::PD-Plug (PD-Plug) and LexA::icals3m (icals3m). Independent factors genotype and treatment were significant, with a significant interaction between the two (ANOVA; F = 82.4, df = 2, P < 0.0001; F = 445.0, df = 1, P < 0.0001; F = 104.2, df = 2, P < 0.0001). Significant differences denoted by a, b and c, p < 0.0001. n = 40. **B)** Starch, sucrose, glucose and fructose content in leaves of 5-week-old plants 72 h post DMSO or estradiol treatment. Independent factors genotype and treatment were significant, with a significant interaction between the two for each metabolite (ANOVA; *starch* F = 4.7, df = 2, P < 0.05, F = 9.2, df = 1, P < 0.05; F = 11.4, df = 2, P < 0.01; *sucrose* F = 26.2, df = 2, P < 0.0001; F = 25.8, df = 1, P < 0.001; F = 26.7, df = 2, P < 0.0001; *glucose* F = 70.0, df = 2, P < 0.0001; F = 74.4, df = 1, P < 0.0001; F = 79.3, df = 2, P < 0.0001; *fructose* F = 28.0, df = 2, P < 0.0001; F = 30.4, df = 1, P < 0.0001; F = 28.8, df = 2, P < 0.0001). Significant differences denoted by a and b, p < 0.01. **C)** Relative leaf area of true leaves 14 days post initial treatment (DMSO or estradiol) in light or shade conditions of Col-0, icals3m, NahG and NahG×LexA::icals3m (NahG×icals3m). Independent factors treatment, genotype and shade condition were significant, with a significant interaction between the pairs and all three (ANOVA; F = 105.5, df = 1, P < 0.0001; F = 84.5, df =3, P < 0.0001; F = 940.8, df = 1, P < 0.0001; F = 100.0, df = 3, P < 0.0001; F = 44.4, df = 1, P < 0.0001; F = 53.9, df = 3, P < 0.0001; F = 53.7, df = 3, P < 0.0001). Significant differences denoted by * p < 0.05, ** p < 0.01, and *** p < 0.001. **D)** Bacterial growth (log_10_ CFU/cm^2^ leaf area) of *Pst* DC3000 3 dpi on uninduced and induced tissue of 4–5-week-old plants. Tissue was treated with DMSO (uninduced) or estradiol (induced) 72 h prior to infection. 4 independent experiments were conducted, with 5 plants per treatment/genotype; N = 20. Independent factors treatment and genotype were significant, with no significant interaction between the two (ANOVA; F = 42.6, df = 1, P < 0.0001; F = 75.8, df = 1, P < 0.0001; F = 0.4, df = 1, P =0.5). Significant differences denoted by a, b and c, p < 0.0005.

Plasmodesmal closure might be expected to restrict photoassimilate distribution between source and sink tissue; a reduction in plasmodesmal permeability and rosette size has been documented in *Arabidopsis* alongside a significant accumulation of soluble sugars and starch (e.g., in the MOVEMENT PROTEIN 17 *MP17* overexpressor, *CHOLINE TRASPORTER-LIKE 1 cher1* mutant, *PDLP5* and *PDLP6* overexpressors; Kronberg et al. 2007; Kraner et al. 2017; Li et al. 2024). Therefore, we quantified sugar accumulation in source tissues following plasmodesmal closure, measuring the sucrose, fructose, glucose, and starch content of source leaves of plants 72 h post treatment. These data reveal a significant increase in all four sugars in estradiol treated LexA::icals3m samples (Fig. 4b). Reasoning that an increase in sugar content might down-regulate photosynthesis, we used chlorophyll fluorescence imaging to infer photosynthetic yield and found there was no difference in photosynthetic yield of PSII (as indicated by Y(II)) after 72 h in Col-0, LexA::PD-Plug and LexA::icals3m between the DMSO or estradiol treatments (Table 1). As there were no changes in the photosynthetic yield that correlated with the increase in sugar content in induced LexA::icals3m, we next hypothesized that plasmodesmal closure and sugar accumulation might activate the expression of sugar transporters and starch biosynthesis genes to reduce the concentration of soluble sugars in the cytosol. However, out of 125 genes known to be sugar and starch related (Supplemental Data Set 10; Büttner and Sauer 2000; Williams et al. 2000; Chen et al. 2010; Monroe and Storm 2018; Abt and Zeeman 2020; Preiser et al. 2020; Smith and Zeeman 2020; David et al. 2022; Bavnhøj et al. 2023), only the transporter *SENESCENCE-ASSOCIATED GENE 29 (SAG29)/SUGARS WILL EVENTUALLY BE EXPORTED TRANSPORTERS 15 (SWEET15)* was significantly upregulated when plasmodesmata are closed, in LexA::icals3m at 48h and 72 h (Supplemental Fig. S19, S20). AT3G20460 was differentially expressed in LexA::icals3m estradiol treatment at 72 h and is proposed to be a monosaccharide transporter (Johnson et al. 2006). However, this gene was downregulated in response to plasmodesmal closure and as the function of this gene is not yet well characterized it is difficult to speculate on any possible role in the sugar/starch phenotype (Johnson et al. 2006). These results indicate that while plasmodesmal closure induces starch and soluble sugar accumulation, it does not perturb photosynthesis or the expression of sugar transporters.

**Table 1.** Photosynthetic yield as indicated by Y(II) following DMSO or estradiol treatment of leaves of Col-0, LexA::PD-Plug (PD-Plug) and LexA::icals3m (icals3m). Pooled average of Y(II) ± SD, with a biological replicate data averaged from 4 leaves per treatment/genotype on a given experimental run where N = 8, combined from two independent experiments. Independent factors genotype and treatment were not significant, with no significant interaction between the two (ANOVA; F = 1.3, df = 2, P = 0.29; F = 1.2, df = 1, P = 0.29, F = 0.1, df = 2, P = 0.92).

|  | Col-0 | PD-Plug | icals3m |
| --- | --- | --- | --- |
| DMSO | 0.79±0.02 | 0.78±0.02 | 0.78±0.02 |
| estradiol | 0.79±0.01 | 0.78±0.01 | 0.79±0.01 |

We reasoned that sugar accumulation might induce osmotic stress and so to test whether sugar accumulation impairs growth and induces senescence in LexA::icals3m, we simultaneously induced plasmodesmal closure and lowered the light intensity by shading the plants to reduce photosynthesis and thus starch and sugar content (Law et al. 2018). We observed that shade conditions reduced growth in all plants, even Col-0, compared to the light grown plants after 14 days of this light regime (Supplemental Fig. S21). However, estradiol treatment of LexA::icals3m had a further size reduction in comparison to the DMSO treatment (Fig. 4c). While senescence of cotyledons was still evident, it was not as severe as that induced in normal light conditions (Supplemental Fig. S21). This suggests that while plasmodesmal closure induces starch and sugar accumulation, this does not impair growth or singularly induce senescence, and that these phenotypes are regulated by other consequences of plasmodesmal closure.

### SA does not mediate plasmodesmal-induced effects on growth and resistance

Given we detected that plasmodesmal closure increases SA and this could induce the observed senescence and reduction in growth in LexA::icals3m, we sought to test if there is a causal relationship. To test the hypothesis that SA accumulation reduces size and/or induces senescence, we crossed LexA::icals3m with plants that constitutively express *NahG*, the bacterial SA hydroxylase that degrades SA (Gaffney et al. 1993; Delaney et al. 1994; Friedrich et al. 1995; NahG×icals3m). We observed a significant reduction in leaf size as well as senescence observed in the cotyledons after 14 days post plasmodesmal closure in both LexA::icals3m and NahG×icals3m plants (Fig. 4c and Supplemental Fig. S21), indicating that SA accumulation is not the cause of these phenotypes. Like LexA::icals3m, NahG×icals3m plants showed reduced growth and mild cotyledon senescence when grown in shade conditions, suggesting that SA does not contribute to the effects of plasmodesmal closure observed in these conditions.

Lastly, we utilized the NahG×icals3m cross to investigate whether plasmodesmal closure enhances pathogen resistance to *Pst* DC3000 by accumulation of SA. We inoculated LexA::icals3m and NahG×icals3m plants 72 h post DMSO or estradiol treatment (Fig. 4d) and found DMSO treated NahG×icals3m had the highest level of bacterial growth 3 days post infection (dpi). However, estradiol treatment of both LexA::icals3m and NahG×icals3m reduced bacterial growth suggesting that SA accumulation is not the primary cause of the increased resistance induced by plasmodesmal closure.

## Discussion

Plants initiate a suite of defense responses when challenged by a microbial threat. Plasmodesmal closure is an early reaction, initiated within 30 minutes of perceiving pathogen-derived molecules and stress-associated signals including SA and hydrogen peroxide (Xu et al. 2017; Cheval et al. 2020; Tee et al. 2023b), and is critical for the execution of a full response. Thus, we reason that plasmodesmal closure regulates a range of downstream processes. However, considering the plethora of responses elicited during immunity, and the array of elicitors that trigger plasmodesmal closure via converging signaling pathways (Tee et al. 2023b), it is challenging to identify what specific contribution plasmodesmal closure makes to the overall execution of immunity. Using lines with which we can induce plasmodesmal closure independently of biotic elicitors, we explored what occurs when plasmodesmata close without other co-incident processes and determined that plasmodesmal closure itself induces stress.

Transcriptional profiling of responses to plasmodesmal closure identified transcriptional signatures of defense responses associated with SA production and signaling. It has long been known that constitutive over-production of the plasmodesmal protein PDLP5 closes plasmodesmata and leads to high levels of SA (Wang et al. 2013). However, as *PDLP5* transcriptionally responds to SA, it is viewed as a stress response element (Wang et al. 2013). We have previously shown that microbial elicitors, flg22 and chitin, also require PDLP5 to elicit plasmodesmal closure (Tee et al. 2023b) posing a potential model in which plasmodesmal closure is first initiated upon pathogen perception, leading to the accumulation of SA at plasmodesmata to create a feedback loop to sustain closure.

The accumulation of SA when plasmodesmata are closed might be elicited directly or indirectly. SA accumulates to higher levels in induced LexA::icals3m (Fig. 2d), in parallel with an increase in sugars (Fig. 4c) and correlated with greater plasmodesmal closure observed in this line relative to LexA::PD-Plug (Fig. 1c). It is possible that the increase in sugar content induces osmotic stress or another process that feeds into SA signaling, and therefore that these stress responses are secondary to plasmodesmal closure. Further, the combined increase in sugar content and SA might explain our pathoassay data; the necrotrophic pathogen *B. cinerea* might be able to access increased intracellular sugar stores and be unimpeded by SA defense pathways (Ferrari et al. 2003; El-Oirdi et al. 2011), while *Pst* DC3000 might be unable to access intracellular sugars and is impeded by SA (Velásquez et al. 2017; Wilson et al. 2017; Howlader et al. 2020). However, as inducing plasmodesmal closure still enhanced resistance to *Pst* DC3000 in the SA deficient NahG×icals3m plants (Fig. 4d), this indicates SA is only part of plasmodesmal stress response for inducing resistance.

Increasingly there is evidence that sugar is critical for coordinating defense in *Arabidopsis*, where specific elevations in glucose-6-phosphate enhance defense responses before a pathogen invasion (Yamada et a. 2024). Sugars can induce specific signal transduction pathways, with some studies finding that increasing source leaf carbohydrate content leads to a feedback loop of decreasing photosynthetic yield (Araya et al. 2006). Additionally, an inverse relationship between plasmodesmal aperture and the expression/activity of sugar transporters has been reported (Ruan et al. 2001; Zhang et al. 2017; Liu et al. 2022) suggesting that cells maintain the capability to transport sugars between cells by balancing plasmodesmal aperture with transporter activity (Zhang et al. 2017). On this basis, we might expect that induction of LexA::icals3m would activate a compensatory mechanism for sugar distribution. However, we only observed that one sugar transporter was upregulated in induced LexA::icals3m, SAG29/SWEET15, which is also activated by SA during the process of leaf senescence (Seo et al. 2011; Wang et al. 2021a). Therefore, while this may be a component of the senescing phenotype observed in induced LexA::icals3m, the increased expression of this transporter does not counteract the intracellular sugar accumulation observed (Chen et al. 2015). Furthermore, when we reduced the SA, sugar and starch content that occurs in LexA::icals3m, impaired growth and senescence still occurred (Fig. 4c and Supplementary Fig. S21). This indicates there is another yet to be determined mechanism that governs these processes in response to plasmodesmal closure. Irrespective, the question arises whether cellular mechanisms that perceive an increase in soluble sugars are incapacitated when plasmodesmata are closed and cells are isolated.

The variance in multiple sets of experimental data suggest that physiological plasmodesmal responses are heterogeneous within a tissue, i.e., not all cells close their plasmodesmata in response to an elicitor such as chitin or flg22 (Faulkner et al. 2013; Xu et al. 2017; Cheval et al. 2020; Tee et al. 2023b). Our transgenic lines show two different patterns of plasmodesmal response: we see variability in GFP mobility in the induced LexA::PD-Plug line that resembles physiological data sets despite homogeneous induction of the transgene (Supplemental Fig. S1), while data from induced LexA::icals3m exhibits much lower variance (Supplemental Fig. S3). This suggests that LexA::icals3m induces plasmodesmal closure more homogeneously than in LexA::PD-Plug, with more cells becoming isolated from their neighbors than occurs physiologically. This greater homogeneity of plasmodesmal closure in LexA::icals3m correlates with the responses measured in most of the assays performed in this study, with effects on disease resistance, growth, ROS burst, sugar and SA accumulation all greater in magnitude in, or only detected in, LexA::icals3m relative to LexA::PD-Plug. Thus, our data suggests that we have detected some of these effects only when they have been elicited in more cells. However, an alternative possibility is that some responses to plasmodesmal closure occur only when a threshold of isolated cells is reached, which would imply that responses to plasmodesmal closure is a tissue level response, and that connectivity is a population-level parameter. Further, that we observe homogeneous cellular isolation can result in undesired outcomes suggests a degree of heterogeneity is optimal for overall physiology.

In this study we have induced plasmodesmal closure to isolate it from other immune responses and study its effects. For this to be a useful pursuit, we needed to determine whether induced plasmodesmal closure mimics physiological plasmodesmal closure. We reasoned that during an infection, the exposure of cells to PAMPs is likely to be a sustained process suggesting that prolonged plasmodesmal closure can occur during infection. This is supported by our findings that plasmodesmal callose deposits were increased 72 h post *Pst hrcC*^-^ infection (Supplemental Fig. S4). Beyond this, we have identified that immune-independent plasmodesmal closure triggers stress-associated responses, raising the question of whether these responses occur when cells are isolated in any context, including developmental transitions. In many investigations of developmental transitions, *icals3m* is used to induce ectopic plasmodesmal closure at different cell boundaries and our data indicates that this is also likely to block the distribution of sugars to, or from, targeted cells and activate defense. Thus, phenotypes that arise from blocking plasmodesmata at a critical developmental transition must be interpreted with these factors in mind.

We have found that plasmodesmal closure triggers stress responses. This suggests that cellular isolation is a stress-inducing state and, conversely, that connection to neighboring cells is essential for optimal growth-associated cellular functions. Thus, understanding how a cell responds when it becomes isolated from its neighbors, and why cell-to-cell connectivity regulates homeostatic cellular processes, is key to understanding how plants function as multicellular organisms.

## Methods

### Plant material

All *Arabidopsis thaliana* used in this study is in the Columbia-0 ecotype background. For assays on mature plants, *Arabidopsis* plants were grown at 10 h light at 22 °C, on soil or for the RNA-seq experiment, on Murashige and Skoog (MS) 1% sucrose 0.8% agar in 50 mm diameter petri dishes for 4–5 weeks. For MAPK and ROS measurement assays on seedlings, seeds were germinated on MS 1% sucrose 0.8% agar at 16 h light at 22 °C and grown for 8 days, then 10 seedlings were transferred per well on a 6-well plate together with 8 mL of liquid MS and grown for an additional 7 days.

### Confocal microscopy

Microprojectile bombardment, transgene induction visualization and aniline blue plasmodesmal callose measurements utilized confocal microscopy (LSM Zeiss 800). RFP/mCherry was excited with a 561-nm DPSS laser and collected at 600 to 640 nm, GFP/H_2_DCFDA was excited with a 488-nm argon laser and collected at 505 to 530 nm, and aniline blue was excited with a 405-nm UV laser and collected at 430 to 550 nm.

### Microprojectile bombardment

Microprojectile bombardment of *35S::GFP* was performed as described in (Tee et al. 2022). 24 h after DMSO or estradiol treatment, 5-week-old leaves were co-bombarded with pB7WG2.0.RFP_ER_ and pB7WG2.0.GFP. Bombardment sites were then imaged 16 h after using confocal microscopy with a 20× water dipping objective (W N-Achroplan 20×/0.5).

### Generation of transgenic lines and induction of transgene

The *LexA::icals3m* construct is as described (Bellandi et al. 2022). The *LexA::PD-Plug* construct was assembled into a binary vector via Golden Gate cloning; first the ‘PD-Plug’ sequence was created by fusing the PDLP1 signal peptide and the PDLP1 transmembrane domain and cytoplasmic tail to the N-and C-termini of mCherry respectively (Caillaud et al. 2014). This synthetic peptide was inserted downstream of the *LexA* promoter sequence and upstream of a NOS terminator and assembled into the backbone of pICSL0022013 containing the *Nos::BAR::Nos* selection cassette and *Act2::XVE::Act2* cassette.

*LexA::PD-Plug* and *LexA::icals3m* constructs were transformed into the Col-0 *Arabidopsis* background and screened with phosphinothricin (10 µg/mL concentration) or kanamycin selection (50 µg/mL concentration) respectively. For induction of the transgene in all experiments (with the exception of size phenotyping), leaves were painted on both the abaxial and adaxial side with either 0.1% DMSO (mock treatment) or 20 µM β-Estradiol 17-acetate (estradiol treatment) containing 0.01% Silwet L-77 in ddH_2_O, using a fine paint brush. For size phenotyping, seedlings were sprayed with the above specified DMSO or estradiol treatment.

### Plasmodesmal callose measurements

24 h post treatment with either DMSO or estradiol, expanded leaves of 5-week-old plants were infiltrated with 1% aniline blue in PBS buffer (pH 7.4) then imaged using confocal microscopy from the abaxial side using a 63× water immersion objective (Plan-Apochromat 63×/1.20). Z-stacks from multiple areas from a given treated leaf, with a minimum of three biological replicates per genotype and treatment were collected. An automated image analysis pipeline to quantify aniline blue-stained plasmodesmata is available at (Johnston et al. 2022).

### Quantitative PCR

To determine transgene expression, *Arabidopsis* T2 seedlings were grown on MS 1% sucrose 0.8% agar with phosphinothricin (10 µg/mL concentration) or kanamycin (50 µg/mL concentration), for 21 days in 16 h light at 22 °C. Leaves from each line was treated with DMSO or estradiol as above and harvested 48 h later in liquid nitrogen. Samples were homogenized in a GenoGrinder 2100, with RNA extracted, DNase treatment, first strand cDNA synthesis and qPCR performed as previously described (Bellandi et al. 2022). Genes tested are *icals3m*/*CalS3* (using primers AGGACGTTACTTCATGGCGG and GGCTCCGGGAATACAGTCTG) and the housekeeper *AtUBQ10* (using primers AGTCTACTCTTCACTTGGTCCTGC and GCCCCAAAACACAAACCACCAAAG). Normalized relative quantities (NRQs) were generated using the qBase model (Hellemans et al. 2008).

### Pseudomonas syringae infection assays

Three leaves from 4–5-week-old plants were treated with DMSO or estradiol. After 72 h, leaves were infiltrated with 5 × 10^4^ cfu/mL *Pseudomonas syringae* DC3000 and harvested 72 h later. Each leaf was harvested with a 5 mm corer and the three leaves from a single plant combined as a sample. Leaf tissue was homogenized with 1 mL ddH_2_O for 2 mins at 1200 rpm in a GenoGrinder 2100. 5 µL of multiple 10-fold dilutions were plated out with three technical replicates per plant sample, then incubated for 48 h at 28 °C with colony counts determined from countable dilutions. Four independent experiments with five plants per treatment/genotype per experiment was performed.

To determine plasmodesmal associated callose deposits after *Pseudomonas syringae* DC3000 *hrcC^-^* (*hrcC^-^*) infection, a culture at an OD of 0.2 was prepared. Leaves from 5-week-old Col-0 plants were infiltrated either with H_2_O (mock) or *hrcC^-^*, and then imaged 24 h, 48 h or 72 h post treatment. Aniline blue staining was performed as per the above plasmodesmal callose measurements with the exception that 0.1% aniline blue was used. 3-4 z-stacks from multiple areas per leaf was performed, with eight biological replicates per treatment/timepoint collected. Only callose deposits at plasmodesmata were analyzed.

### Botrytis cinerea infection assays

*B. cinerea* spores were applied to leaves from 4–5-week-old plants 72 h post DMSO or estradiol treatment. *B. cinerea* spores were harvested and adjusted to a concentration of 2.5 × 10^5^ spores/mL in 0.25× potato dextrose broth and incubated at room temperature for 2 h with continual shaking for spore germination. Leaves were adhered to 1.5% agar H_2_O plates, and droplets of 2 µL spore suspensions were placed in between the mid vein and leaf edge on each side. Plates were sealed with parafilm and incubated in a cabinet set at 22 °C with 10 h light. Developing disease lesions were photographed 2 dpi and measured using FIJI. Each prepared plate had four leaves per genotype and treatment, and the average lesion area was calculated per genotype/treatment in a given plate, defined as a single biological replicate. Three independent experiments were conducted with a minimum of five biological replicates per genotype/treatment.

### MAPK activation assays

MAPK activation assays were performed on leaf discs from expanded leaves of 5-week-old plants or on 15-day-old seedlings 72 h post treatment with DMSO or estradiol. Samples were harvested 0, 5 mins, 15 mins, 30 mins and 45 mins post 100 nM flg22 treatment, and flash frozen in liquid nitrogen. Frozen samples were homogenized via 90 seconds of shaking (1100 rpm) in a GenoGrinder 2100, then 500 µL protein extraction buffer (50 mM Tris-HCl pH 7.5, 150 mM NaCl, 1 mM EDTA pH 8.0, 10% glycerol, 0.5% NP40 IGEPAL® CA-630, protease inhibitor cocktail (Sigma) 1:100, phosphatase inhibitor (Sigma) 1:200, 1 mM Na_2_MoO_4_×2H_2_O, 1mM NaF, 1.5 mM activated Na_3_VO_4_, 5 mM DTT, 1mM PMSF) was added and the sample left to mix for 30 mins at 4 °C. Samples were centrifuged at max speed for 10 mins at 4 °C, supernatant transferred to a new tube and re-centrifuged at max speed for another 10 mins at 4 °C. The resulting supernatant was incubated at 95 °C for 15 mins with Laemmli buffer, then proteins were separated by SDS-PAGE and transferred to Immuno-blot® PVDF membrane. Proteins were detected using p44/42 MAPK (1:2000; Cell Signaling Technologies, #9102) and α-Rabbit-HRP (1:10000, Sigma, A0545).

### ROS burst measurements

ROS assays were performed on leaf discs taken from 5-week-old plants or 15-day-old seedlings 72 h post DMSO or estradiol treatment. Leaf discs were first floated on H_2_O overnight; the following day water was removed, then 20 µg/mL HRP and 6 µM L-012 was added, with or without 100 nm flg22. The same was performed for seedlings except liquid media was removed before adding the HRP and L-012. Chemiluminescence was recorded using a Varioskan Flash (Thermo Fisher) over 45 mins, with the total luminescence emitted used for analysis.

### General ROS quantification

20 µM H_2_DCFDA (Thermo Fisher) was syringe infiltrated into expanded leaves of 5-week-old plants 72 h post DMSO or estradiol treatment, and confocal microscopy was performed on the abaxial side of the leaf. Z-stacks were taken from four areas of a given leaf from four plants per genotype/treatment. Quantification of fluorescence was performed in FIJI using a sum projection.

### RNA-seq

Two mature leaves per 5-week-old plant were DMSO or estradiol treated, and harvested 12 h, 24 h, 48 h and 72 h post treatment. Each biological replicate contained four plants (i.e., 8 leaves), with three biological replicates for each genotype/treatment/time point. Leaves were flash frozen in liquid nitrogen, then homogenized via 90 seconds of shaking (1100 rpm) in a 2100 Geno/Grinder (Spex SamplePrep, USA). Total RNA was extracted using the RNEasy Plant Mini Kit (QIAGEN, Germany) and eluted into 60 µl of water. Purified RNA was treated with the TURBO DNA-free kit (Thermo Fisher, USA). RNA quantification and library construction was conducted by Novogene Co., Ltd (Beijing). RNA quality and quantity were access via 1% agarose gel, NanoPhotometer ® spectrophotometer (IMPLEN, USA), and the RNA Nano 6000 Assay Kit for the Bioanalyzer 2100 System (Agilent Technologies, USA). Libraries were constructed using 0.4 µg of RNA and the NEBNext ® UltraTM RNA Library Prep Kity for Illumina (NEB, USA). mRNA was purified via poly-T magnetic beads. Fragmentation, cDNA synthesis, and NEBNext Adaptor hybridization were performed according to manufacturer’s instructions and purified with the AMPure XP system (Beckman Coulter, USA). Index-coded samples were clustered on a cBot Cluster Generation System using TruSeq PE Cluster Kit v3-cBot-HS (Illumina, USA) and sequenced on an Illumina Novaseq platform to generate 150bp paired-end reads with a sequencing depth of 10 million reads per sample. All sequencing data are available on GEO at GSE248301.

### Bioinformatic analyses

Adaptors and low-quality (phred33<20) reads were trimmed from the data using TrimGalore v0.5.0 (Krueger et al. 2021) and Cutadapt v1.7 (Martin 2011). RNA quality was accessed using FastQC v0.11.8 (Babraham Bioinformatics - FastQC A Quality Control tool for High Throughput Sequence Data) and MultiQC v.1.7 (Ewels et al. 2016). The data was mapped against the *Arabidopsis* reference genome (TAIR10; NCBI Refseq GCA_000001735.1) using HISAT2 v2.1.0 (Kim et al. 2019). Mapped reads were sorted using Samtools v1.4.1 Sort (Danecek et al. 2021). Transcript assembly was completed using Stringtie v1.3.3 (Pertea et al. 2015) and gene counts (see Supplemental Data Set 3 and 11 for gene count matrix and metadata respectively) were extracted using the built-in python wrapper, prepDE.py available at http://ccb.jhu.edu/software/stringtie/dl/prepDE.py. Principle component analysis was conducted using DEseq2, ggplot2 (Wickham 2016), ggConvexHull, and TidyVerse (Wickham et al. 2019) in R (R Core Team 2022). Differentially expressed genes were determined using DEseq2 (Love et al. 2014) in R. Genes with less than 10 counts across all samples were excluded from the analysis. The data was subset by timepoints, and Wald Tests or Likelihood Ratio Tests (LRT) were performed based on comparisons of interest. Wald Tests were used to extract DEGs based on differences between treatment or genotype. LRT were used to extract genes differentially expressed based on difference between closed plasmodesmata (induced/estradiol treated LexA::PD-Plug and LexA::icals3m combined) and open plasmodesmata (uninduced/DMSO treated LexA::PD-Plug, uninduced/DMSO treated LexA::icals3m, and DMSO and estradiol treated Col-0). LRT was used in this way to determine genes uniquely up- or down-regulated in the inducible lines treated with estradiol. Differentially expressed genes from the Wald Test were designated if the log2 fold change (lfc) was greater than |1.5| and adjusted p-value<0.05. Differentially expressed genes from the LRT were considered significant if the adjusted p-value was less than 0.05. LRT significantly differentially expressed genes were clustered using Lasso2 (Lokhorst et al. 2021) and DEGreports v1.36.0 (Pantano 2023) and normalized read counts plotted using ggplot2 and pheatmaps v1.0.12 (Kolde 2019). Genes involved in Salicylic Acid biosynthesis/catabolism and response were determined based on a Wald Test of open plasmodesmata versus closed plasmodesmata with a lfc cut-off of +/-0.5 and adjusted p-value of 0.05 (Supplemental Data Set 12). Gene ontology enrichment analysis was conducted based on differentially expressed genes. Enrichment was conducted in TopGo v2.52.0 (Alexa and Rahnenfuhrer 2023) based on TAIR10 GO terms, with a topgoFisher cut-off of p<0.05. GO terms were plotted in ggplot2. GO parent terms were determined using the Revigo’s (Supek et al. 2011) redundancy feature (cut-off 0.7) and utilized to extract defense-specific enriched GO terms.

### Transgene Mapping

Transgenes were mapped by generating a pseudo-genome consisting of the *Arabidopsis* TAIR genome as well as the transgene of interest (mCherry) using SeqKit v0.9.1 (Shen et al. 2016) and STAR v2.5a (Dobin et al. 2013). The reads were then mapped and sorted by coordinates to the pseudo-genome using STAR v2.5a. The data was processed the same as previously described using Samtools and Stringtie. Gene counts of mCherry (Supplemental Data Set 13) and CalS3 (Supplemental Data Set 14) were extracted using DESeq2 and plotted in ggplot2.

### Size phenotyping

9-day-old seedlings of Col-0, LexA::PD-Plug and LexA:: icals3m were finely sprayed with treatments DMSO or estradiol and 6 days later, the first two true leaves were adhered to a microscope slide and images taken on a Leica M205 FA stereo microscope. Leaf area was measured using FIJI. For the shade experiments, 10-day-old seedlings of Col-0, LexA::icals3m, NahG and NahG×icals3m were finely sprayed with treatments of DMSO or estradiol and either kept at normal light conditions (80 μmol/m^-2^/s^-1^) or a shaded lid was placed on top reducing the light intensity to 9 μmol/m^-2^/s^-1^. An additional spray was performed 7 days post the initial treatment, and 14 days post treatment the first two true leaves were adhered to a microscope slide and images taken on a Axio Zoom V16 light microscope. A leaf measurement macro developed using FIJI was used to measure the leaf area; the average between the two true leaves per plant was calculated, and the relative size was calculated by dividing the average of the DMSO control in each condition/genotype for each sample.

### Sugar and starch measurements

Leaf tissue was harvested in liquid nitrogen from plants 72 h post treatment of either DMSO or estradiol (3 leaves from 5-week-old plants), with sugars and starch separately extracted from approximately 100 mg total leaf tissue as described by (Smith and Zeeman 2006). Frozen tissue was homogenized for 2 mins at 1200 rpm in a GenoGrinder 2100. Samples were further homogenized in 1 mL of 0.7 M cold perchloric acid and then centrifuge at max speed 4 °C for 5 min. The supernatant was transferred to process for sugar, while the pellet was kept on ice for starch quantification. For sugar, 600 µL supernatant was transferred to a new 1.5 mL tube, and neutralization buffer (2 M KOH, 400 mM MES) was added to achieve a pH of 6-7. The sample was then spun at max speed for 4 min and supernatant recovered. Soluble sugars were quantified by enzymatic assay in technical triplicates by monitoring NADH production absorbance at 340 nm in a 96-well format using an Omega FLUOstar microplate reader. Each sample was added to 50 µL of buffer (50 mM HEPES NaOH pH 7.4-7.6, 1 mM MgCl_2_, 1mM ATP, 1 mM NAD, 1.4 U Hexokinase), topped with H_2_O to a total volume of 198 µL. An initial A_340_ reading was taken, before adding glucose-6-phosphate dehydrogenase for a kinetic reading to determine glucose content, followed by adding phosphoglucose isomerase to determine fructose content, followed by invertase to determine sucrose content. To determine starch content, the sample pellet was first washed by adding 1 mL H_2_O and vortexing, followed by centrifugation for 3 mins at 3,000 ×g. Supernatant was removed, then pellet washed again by adding 1 mL 80% ethanol and vortexing, followed by centrifugation for 3 mins at 3,000 ×g; this ethanol wash was repeated for a total of three washes. Excess ethanol was evaporated, then resuspended with 750 µL of H_2_O. 2 × 200 µL of each sample was transferred to a new screw cap tube and incubated at 95 °C for 12 minutes. The samples were left to briefly cool to room temperature (∼5 min), then one aliquot was digested by incubation with α-amylase (10 U) and amyloglucosidase (1.26 U) in 0.1045 M sodium acetate buffer (pH 4.8) for two hours at 37 °C, alongside the other aliquot serving as the non-digested control (i.e., containing only the 0.1045 M sodium acetate buffer and two hour 37 °C incubation). Starch content was quantified through enzymatic assay of the digested aliquot sample by monitoring NADH production as per the above glucose measurement. Technical triplicates were performed for each digested sample, alongside a single technical replicate for the non-digested sample. Starch content was calculated from the glucose assay values in mg per gram of fresh weight of sample.

### Hormone measurements

Leaves from 5-week-old plants were treated with DMSO or estradiol and harvested 72 h post treatment and flash frozen in liquid nitrogen; per biological replicate, a minimum of six leaves from a minimum of three plants were pooled together to obtain between 150-220mg of fresh tissue weight. Frozen tissue was homogenized for 2 × 2 mins at 1200 rpm in a GenoGrinder 2100. 300 µL of buffer (10% methanol, 1% acetic acid in H_2_O) containing an internal standard of 1 µM salicylic acid-d_4_ (SA-_d4_, Merck) and 500 nM jasmonic acid-d_5_ (JA_-d5_, Cayman Chemicals) was added, mixed well with a vortex then left to mix at 4 °C for 20 mins. Samples were centrifuged for 25 mins at 4°C at 15,000 RCF, and the supernatant recovered. The pellet was re-extracted with another 300 µL buffer without the internal standard, mixed and centrifuged as before, with the resulting supernatant combined with the first recovered supernatant. The total supernatant was centrifuged at 4 °C for 10 mins at max speed, and each sample was then run for analysis. Quantification was performed on an Acquity UPLC attached to a Xevo TQS tandem mass spectrometer (Waters). Separation was on a 50×2.1mm 2.6µM Kinetex EVO C18 column (Phenomenex) using the following gradient of acetonitrile (solvent B) versus 0.1% formic acid in water (solvent A), run at 600 µL.min-1 and 35°C: 0 min, 10% B; 2 min, 80% B; 2.15 min, 80% B; 2.2 min, 10% B; 3.4 min, 10% B. The injection volume was 5 µL. Salicylate was quantified by negative mode electrospray, monitoring the transition *m/z* 137→65 (and for the internal standard the corresponding transition 141→97) while jasmonate was monitored at the transition *mz* 209→59 (and for the internal standard the corresponding transition 214→62) at 16V collision energy. Spray chamber conditions were 1.5kV spray voltage, 500°C desolvation temperature, 900 L.hr^-1^ desolvation gas, 150 L.hr^-1^ cone gas, and 7 bar nebulizer pressure. Results were processed using TargetLynx software V4.1 SCN876 (2012).

### Chlorophyll fluorescence measurements

Chlorophyll fluorescence measurements were performed using an Imaging-PAM chlorophyll fluorometer and ImagingWin software application (Walz). The effective PSII quantum yield as indicated by Y(II) was taken following a 30 min dark adaptation of leaves from 5-week-old plants 72 h post DMSO or estradiol treatment. Four leaves per treatment/genotype were measured in a given run, with 8 runs in total over two independent experiments.

### Statistical analyses

Statistical analyses were performed using RStudio 2021.09.01 Build 351/R version 4.0.3. Data presented as boxplots have the middle line indicating the median, the box representing the upper and lower quartiles, and the whiskers indicating the minimum and maximum values with 1.5× interquartile range. For bootstrap analyses, data were analyzed using *medianBootstrap* (Johnston and Faulkner 2021) or *medianBootstrap2* (Tee et al. 2023a). Pathoassays, basal ROS quantification, and chlorophyll fluorescence measurements were analyzed using a linear-mixed effects model using the R package, *lmerTest*. For pathoassays the random effect was ‘independent experiment repeat’; for the basal ROS quantification the random effect was ‘biological replicate’; for chlorophyll fluorescence measurements the random effect was ‘experimental run’. Size phenotyping, starch, sucrose, glucose and fructose measurements were analyzed using a linear model. All ANOVAs specified are ANOVA Satterthwaite’s Method, and significant differences between factors were determined by post hoc Tukey HSD using the R package, *emmeans*. Salicylic acid quantification was analyzed using a Mann-Whitney test.

## Data availability

Raw counts for bombardment data generated for this study (Fig. 1C and Supplemental Fig. S3) is available in Supplemental Data Set 1. Additional raw counts for bombardment data for Supplemental Fig. S3 are sourced from Cheval et al. 2020 and Tee et al. 2023b. RNAseq data has been deposited in the Gene Expression Omnibus under the accession code GSE248301, with further analysis included in Supplemental Data Sets as specified in-text.

## Supplemental Data Sets

Supplemental Data Set 1: Data for number of cells showing GFP within a genotype/treatment

Supplemental Data Set 2: Background response to estradiol: differentially regulated genes in response to estradiol treatment shared across all genotypes

Supplemental Data Set 3: Gene count matrix from the RNAseq analysis

Supplemental Data Set 4: Genes differentially expressed in category specified, identified by a likelihood ratio test analysis

Supplemental Data Set 5: Genes differentially expressed in response to 12h plasmodesmal closure identified by a likelihood ratio test analysis

Supplemental Data Set 6: Genes differentially expressed in response to 24h plasmodesmal closure identified by a likelihood ratio test analysis

Supplemental Data Set 7: Genes differentially expressed in response to 48 h plasmodesmal closure identified by a likelihood ratio test analysis

Supplemental Data Set 8: Genes differentially expressed in response to 72h plasmodesmal closure identified by a likelihood ratio test analysis

Supplemental Data Set 9: Genes identified from the literature as associated with salicylic acid biosynthesis, catabolism and response

Supplemental Data Set 10: Genes identified from the literature as associated with the synthesis and transport of soluble sugars and starch

Supplemental Data Set 11: Metadata related to the gene count matrix

Supplemental Data Set 12: Log2 fold change of genes associated with SA that are differentially expressed at 12h, 24h, 48h and 72h

Supplemental Data Set 13: Gene counts of mCherry from pseudo-genome GEO GSE248301

Supplemental Data Set 14: Gene counts of CalS3 from pseudo-genome GEO GSE248301

## Author Contributions

CREDIT authorship breakdown:

Conceptualization – all authors Data curation – EET, ABr, DP, HRT

Formal analysis – EET, ABr, DP, HRT

Funding acquisition – CF

Investigation – EET, ABr, DP, HRT, CW, Abe

Methodology – EET, ABr, DP, HRT, CW, ABe, CF

Project administration – CF

Resources – DP, ABr, CF

Supervision – CF

Validation – EET, ABr, DP, HRT

Visualization – EET, ABr, DP, HRT

Writing – original draft – EET, HRT, CF

Writing – review & editing – EET, ABr, DP, HRT, CW, ABe, CF

## Acknowledgements

We acknowledge access to the John Innes Centre Biomolecular Analysis Facility and thank Dr. Lionel Hill for their assistance and training with mass spectrometry, as well as access to the John Innes Centre Bioimaging Facility and thank Dr Sergio G Lopez for support with imaging. We acknowledge the Seung lab for technical assistance in the sugar and starch measurements. We would like to thank Qi Yang Ngai for their assistance in developing the leaf measurement macro and Núria Real-Tortosa for critical reading of the manuscript. The *NahG* seeds were kindly provided by Jurriann Ton (The University of Sheffield).

Work in the Faulkner lab is funded by the Biotechnology and Biological Science Research Council (Grants: BB/X010996/1 and BB/X007685/1) and the European Research Council (Grant: 725459 - “INTERCELLAR).

## References

1. Abt MR and Zeeman SC. Evolutionary innovations in starch metabolism. Curr Opin Plant Biol. 2020:55:109–117. 10.1016/J.PBI.2020.03.001

2. Alazem M and Burch-Smith TM. Roles of ROS and redox in regulating cell-to-cell communication: Spotlight on viral modulation of redox for local spread. Plant Cell Environ. 2024. 10.1111/PCE.14805

3. Alexa A and Rahnenfuhrer J. topGO: Enrichment Analysis for Gene Ontology. 2023.

3a. Araya T, Noguchi K, and Terashima I. Effects of Carbohydrate Accumulation on Photosynthesis Differ between Sink and Source Leaves of Phaseolus vulgaris L. Plant Cell Physiol. 2006:47(5):644–652. 10.1093/PCP/PCJ033

4. Aung K, Kim P, Li Z, Joe A, Kvitko B, Alfano JR. and He SY. 2020. Pathogenic bacteria target plant plasmodesmata to colonize and invade surrounding tissues. The Plant Cell, 32(3), pp.595–611. 10.1105/tpc.19.00707

5. Babraham Bioinformatics - FastQC A Quality Control tool for High Throughput Sequence Data. https://www.bioinformatics.babraham.ac.uk/projects/fastqc/. Retrieved July 28, 2023

6. Bavnhøj L, Driller JH, Zuzic L, Stange AD, Schiøtt B, and Pedersen BP. Structure and sucrose binding mechanism of the plant SUC1 sucrose transporter. Nature Plants 2023 9:6. 2023:9(6):938–950. 10.1038/s41477-023-01421-0

7. Beaugelin I, Chevalier A, d’Alessandro S, Ksas B, and Havaux M. (2020). Endoplasmic reticulum-mediated unfolded protein response is an integral part of singlet oxygen signalling in plants. The Plant Journal, 102(6), 1266–1280. 10.1111/tpj.14700.

8. Bellandi A, Papp D, Breakspear A, Joyce J, Johnston MG, de Keijzer J, Raven EC, Ohtsu M, Vincent TR, Miller AJ, et al. Diffusion and bulk flow of amino acids mediate calcium waves in plants. Sci Adv. 2022:8(42):6693. DOI: 10.1126/sciadv.abo6693.

9. van Butselaar T and Van den Ackerveken G. Salicylic Acid Steers the Growth– Immunity Tradeoff. Trends Plant Sci. 2020:25(6):566–576. 10.1016/J.TPLANTS.2020.02.002

10. Büttner M and Sauer N. Monosaccharide transporters in plants: structure, function and physiology. Biochimica et Biophysica Acta (BBA) - Biomembranes. 2000:1465(1–2):263–274. 10.1016/S0005-2736(00)00143-7

11. Caillaud MC, Wirthmueller L, Sklenar J, Findlay K, Piquerez SJM, Jones AME, Robatzek S, Jones JDG, and Faulkner C. The Plasmodesmal Protein PDLP1 Localises to Haustoria-Associated Membranes during Downy Mildew Infection and Regulates Callose Deposition. PLoS Pathog. 2014:10(11):e1004496. 10.1371/JOURNAL.PPAT.1004496

12. Chen LQ, Hou BH, Lalonde S, Takanaga H, Hartung ML, Qu XQ, Guo WJ, Kim JG, Underwood W, Chaudhuri B, et al. Sugar transporters for intercellular exchange and nutrition of pathogens. Nature 2010 468:7323. 2010:468(7323):527–532. 10.1038/nature09606

13. Chen LQ, Lin IW, Qu XQ, Sosso D, McFarlane HE, Londoño A, Samuels AL, and Frommer WB. A Cascade of Sequentially Expressed Sucrose Transporters in the Seed Coat and Endosperm Provides Nutrition for the Arabidopsis Embryo. Plant Cell. 2015:27(3):607–619. 10.1105/TPC.114.134585

14. Cheval C, Samwald S, Johnston MG, Keijzer J de, Breakspear A, Liu X, Bellandi A, Kadota Y, Zipfel C, and Faulkner C. Chitin perception in plasmodesmata characterizes submembrane immune-signaling specificity in plants. Proc Natl Acad Sci U S A. 2020:117(17):9621–9629. 10.1073/PNAS.1907799117

15. Danecek P, Bonfield JK, Liddle J, Marshall J, Ohan V, Pollard MO, Whitwham A, Keane T, McCarthy SA, and Davies RM. Twelve years of SAMtools and BCFtools. Gigascience. 2021:10(2):1–4. 10.1093/GIGASCIENCE/GIAB008

16. David LC, Lee SK, Bruderer E, Abt MR, Fischer-Stettler M, Tschopp MA, Solhaug EM, Sanchez K, and Zeeman SC. BETA-AMYLASE9 is a plastidial nonenzymatic regulator of leaf starch degradation. Plant Physiol. 2022:188(1):191–207. 10.1093/PLPHYS/KIAB468

17. Delaney TP, Uknes S, Vernooij B, Friedrich L, Weymann K, Negrotto D, Gaffney T, Gut-Rella M, Kessmann H, Ward E, et al. A central role of salicylic acid in plant disease resistance. Science. 1994:266(5188):1247–1250. 10.1126/SCIENCE.266.5188.1247

18. Dobin A, Davis CA, Schlesinger F, Drenkow J, Zaleski C, Jha S, Batut P, Chaisson M, and Gingeras TR. STAR: ultrafast universal RNA-seq aligner. Bioinformatics. 2013:29(1):15–21. 10.1093/BIOINFORMATICS/BTS635

19. El-Oirdi M, El-Rahman TA, Rigano L, El-Hadrami A, Rodriguez MC, Daayf F, Vojnov A, and Bouarab K. Botrytis cinerea Manipulates the Antagonistic Effects between Immune Pathways to Promote Disease Development in Tomato. Plant Cell. 2011:23(6):2405. 10.1105/TPC.111.083394

20. Ewels P, Magnusson M, Lundin S, and Käller M. MultiQC: summarize analysis results for multiple tools and samples in a single report. Bioinformatics. 2016:32(19):3047–3048. 10.1093/BIOINFORMATICS/BTW354

21. Faulkner C, Petutschnig E, Benitez-Alfonso Y, Beck M, Robatzek S, Lipka V, and Maule AJ. LYM2-dependent chitin perception limits molecular flux via plasmodesmata. Proc Natl Acad Sci U S A. 2013:110(22):9166–9170. 10.1073/PNAS.1203458110/-/DCSUPPLEMENTAL

22. Ferrari S, Plotnikova JM, De Lorenzo G, and Ausubel FM. Arabidopsis local resistance to Botrytis cinerea involves salicylic acid and camalexin and requires EDS4 and PAD2, but not SID2, EDS5 or PAD4. The Plant Journal. 2003:35(2):193–205. 10.1046/J.1365-313X.2003.01794.X

23. Friedrich L, Vernooij B, Gaffney T, Morse A, and Ryals J. Characterization of tobacco plants expressing a bacterial salicylate hydroxylase gene. Plant Mol Biol. 1995:29(5):959–968. 10.1007/BF00014969/METRICS

24. Gaffney T, Friedrich L, Vernooij B, Negrotto D, Nye G, Uknes S, Ward E, Kessmann H, and Ryals J. Requirement of Salicylic Acid for the Induction of Systemic Acquired Resistance. Science. 1993:261(5122):754–756. 10.1126/SCIENCE.261.5122.754

25. German L, Yeshvekar R, and Benitez-Alfonso Y. Callose metabolism and the regulation of cell walls and plasmodesmata during plant mutualistic and pathogenic interactions. Plant Cell Environ. 2023:46(2):391–404. 10.1111/PCE.14510

26. Hellemans J, Mortier G, De Paepe A, Speleman F, and Vandesompele J. qBase relative quantification framework and software for management and automated analysis of real-time quantitative PCR data. Genome Biol. 2008:8(2):1–14. 10.1186/GB-2007-8-2-R19/COMMENTS

27. Howlader P, Bose SK, Jia X, Zhang C, Wang W, and Yin H. Oligogalacturonides induce resistance in Arabidopsis thaliana by triggering salicylic acid and jasmonic acid pathways against Pst DC3000. Int J Biol Macromol. 2020:164:4054–4064. 10.1016/J.IJBIOMAC.2020.09.026

28. Johnson DA, Hill JP, and Thomas MA. The monosaccharide transporter gene family in land plants is ancient and shows differential subfamily expression and expansion across lineages. BMC Evol Biol. 2006:6(1):1–20. 10.1186/1471-2148-6-64/FIGURES/10

29. Johnston MG, Bellandi A, and Faulkner C. faulknerfalcons/PD_detection: Toolkit for quantification of callose at plasmodesmata. 2022. 10.5281/ZENODO.6583765

30. Johnston MG and Faulkner C. A bootstrap approach is a superior statistical method for the comparison of non-normal data with differing variances. New Phytologist. 2021:230(1):23–26. 10.1111/NPH.17159

31. Kim D, Paggi JM, Park C, Bennett C, and Salzberg SL. Graph-based genome alignment and genotyping with HISAT2 and HISAT-genotype. Nat Biotechnol. 2019:37(8):907–915. 10.1038/S41587-019-0201-4

32. Kolde R. Package “pheatmap.” 2019.

33. Kraner ME, Link K, Melzer M, Ekici AB, Uebe S, Tarazona P, Feussner I, Hofmann J, and Sonnewald U. Choline transporter-like1 (CHER1) is crucial for plasmodesmata maturation in Arabidopsis thaliana. The Plant Journal. 2017:89(2):394–406. 10.1111/TPJ.13393

34. Kronberg K, Vogel F, Rutten T, Hajirezaei MR, Sonnewald U, and Hofius D. The Silver Lining of a Viral Agent: Increasing Seed Yield and Harvest Index in Arabidopsis by Ectopic Expression of the Potato Leaf Roll Virus Movement Protein. Plant Physiol. 2007:145(3):905–918. 10.1104/PP.107.102806

35. Krueger F, James F, Ewels P, Afyounian E, and Schuster-Boeckler B. FelixKrueger/TrimGalore: v0.6.7 - DOI via Zenodo. 2021. 10.5281/ZENODO.5127899

36. Law SR, Chrobok D, Juvany M, Delhomme N, Lindén P, Brouwer B, Ahad A, Moritz T, Jansson BS, Gardeström P, et al. Darkened Leaves Use Different Metabolic Strategies for Senescence and Survival. Plant Physiol. 2018:177(1):132–150. 10.1104/PP.18.00062

37. Lee J-Y, Wang X, Cui W, Sager R, Modla S, Czymmek K, Zybaliov B, Wijk K van, Zhang C, Lu H, et al. A Plasmodesmata-Localized Protein Mediates Crosstalk between Cell-to-Cell Communication and Innate Immunity in Arabidopsis. Plant Cell. 2011:23(9):3353–3373. 10.1105/TPC.111.087742

38. Levy A, Erlanger M, Rosenthal M, and Epel BL. A plasmodesmata-associated beta-1,3-glucanase in Arabidopsis. Plant J. 2007:49(4):669–682. 10.1111/J.1365-313X.2006.02986.X

39. Li Z, Variz H, Chen Y, Liu SL, Aung K. Plasmodesmata-Dependent Intercellular Movement of Bacterial Effectors. Front Plant Sci. 2021 Mar 22;12:640277. doi: 10.3389/fpls.2021.640277. PMID: 33959138; PMCID: PMC8095247.

40. Li Z, Liu SL, Montes-Serey C, Walley JW, Aung K. PLASMODESMATA-LOCATED PROTEIN 6 regulates plasmodesmal function in Arabidopsis vasculature. Plant Cell. 2024 Jun 6:koae166. doi: 10.1093/plcell/koae166. Epub ahead of print. PMID: 38842334.

41. Liu YH, Song YH, and Ruan YL. Sugar conundrum in plant–pathogen interactions: roles of invertase and sugar transporters depend on pathosystems. J Exp Bot. 2022:73(7):1910. 10.1093/JXB/ERAB562

42. Lokhorst J, Venables B, and Turlach B. R package “lasso2.” 2021. https://github.com/cran/lasso2. Retrieved July 28, 2023

43. Love MI, Huber W, and Anders S. Moderated estimation of fold change and dispersion for RNA-seq data with DESeq2. Genome Biol. 2014:15(12):1–21. 10.1186/S13059-014-0550-8/FIGURES/9

44. Martin M. Cutadapt removes adapter sequences from high-throughput sequencing reads. EMBnet J. 2011:17(1):10–12. 10.14806/EJ.17.1.200

45. Monroe JD and Storm AR. Review: The Arabidopsis β-amylase (BAM) gene family: Diversity of form and function. Plant Science. 2018:276:163–170. 10.1016/J.PLANTSCI.2018.08.016

46. Ohtsu M, Jennings J, Johnston M, Breakspear A, Liu X, Stark K, Morris RJ, de Keijzer J, Faulkner C. Assaying Effector Cell-to-Cell Mobility in Plant Tissues Identifies Hypermobility and Indirect Manipulation of Plasmodesmata. Mol Plant Microbe Interact. 2024 Feb;37(2):84–92. doi: 10.1094/MPMI-05-23-0052-TA. Epub 2024 Feb 26. PMID: 37942798.

47. Pantano L. DEGreport: Report of DEG analysis. 2023. 10.18129/B9.bioc.DEGreport

48. Paterlini A, Dorussen D, Fichtner F, van Rongen M, Delacruz R, Vojnović A, Helariutta Y and Leyser O. (2021), Callose accumulation in specific phloem cell types reduces axillary bud growth in *Arabidopsis thaliana*. New Phytol, 231: 516–523. 10.1111/nph.17398

49. Pertea M, Pertea GM, Antonescu CM, Chang TC, Mendell JT, and Salzberg SL. StringTie enables improved reconstruction of a transcriptome from RNA-seq reads. Nature Biotechnology 2015 33:3. 2015:33(3):290–295. 10.1038/nbt.3122

50. Preiser AL, Banerjee A, Weise SE, Renna L, Brandizzi F, and Sharkey TD. Phosphoglucoisomerase Is an Important Regulatory Enzyme in Partitioning Carbon out of the Calvin-Benson Cycle. Front Plant Sci. 2020:11:580726. 10.3389/FPLS.2020.580726/BIBTEX

51. R Core Team. R: A language and environment for statistical computing. R Foundation for Statistical Computing, Vienna, Austria. 2022. https://www.R-project.org/

52. Ross-Elliott TJ, Jensen KH, Haaning KS, Wager BM, Knoblauch J, Howell AH, Mullendore DL, Monteith AG, Paultre D, Yan D, Otero S, Bourdon M, Sager R, Lee JY, Helariutta Y, Knoblauch M, Oparka KJ (2017) Phloem unloading in Arabidopsis roots is convective and regulated by the phloem-pole pericycle eLife 6:e24125 10.7554/eLife.24125

53. Ruan YL, Llewellyn DJ, and Furbank RT. The Control of Single-Celled Cotton Fiber Elongation by Developmentally Reversible Gating of Plasmodesmata and Coordinated Expression of Sucrose and K+ Transporters and Expansin. Plant Cell. 2001:13(1):47–60. 10.1105/TPC.13.1.47

54. Seo PJ, Park JM, Kang SK, Kim SG, and Park CM. An Arabidopsis senescence-associated protein SAG29 regulates cell viability under high salinity. Planta. 2011:233(1):189–200. 10.1007/S00425-010-1293-8

55. Sevilem I, Miyashima S, and Helariutta Y. (2012). Cell-to-cell communication via plasmodesmata in vascular plants. Cell Adhesion & Migration. 7(1), 27–32. 10.4161/cam.22126

56. Seyfferth C, Tsuda K, Lu H, and Grant M. Salicylic acid signal transduction: the initiation of biosynthesis, perception and transcriptional reprogramming. 2014. 10.3389/fpls.2014.00697

57. Shen W, Le S, Li Y, and Hu F. SeqKit: A Cross-Platform and Ultrafast Toolkit for FASTA/Q File Manipulation. PLoS One. 2016:11(10):e0163962. 10.1371/JOURNAL.PONE.0163962

58. Smith AM and Zeeman SC. Quantification of starch in plant tissues. Nature Protocols 2006 1:3. 2006:1(3):1342–1345. 10.1038/nprot.2006.232

59. Smith AM and Zeeman SC. Starch: A Flexible, Adaptable Carbon Store Coupled to Plant Growth. Annual Review of Plant Biology. 2020:71:217–245. 10.1146/ANNUREV-ARPLANT-050718-100241

60. Spoel SH, Koornneef A, Claessens SMC, Korzelius JP, Van Pelt JA, Mueller MJ, Buchala AJ, Métraux JP, Brown R, Kazan K, et al. NPR1 Modulates Cross-Talk between Salicylate- and Jasmonate-Dependent Defense Pathways through a Novel Function in the Cytosol. Plant Cell. 2003:15(3):760–770. 10.1105/TPC.009159

61. Supek F, Bošnjak M, Škunca N, and Šmuc T. REVIGO Summarizes and Visualizes Long Lists of Gene Ontology Terms. PLoS One. 2011:6(7):e21800. 10.1371/JOURNAL.PONE.0021800

62. Tee EE, Johnston MG, and Faulkner C. faulknerfalcons/Tee-etal-2023. 2023a. 10.5281/ZENODO.7680790

63. Tee EE, Johnston MG, Papp D, and Faulkner C. A PDLP-NHL3 complex integrates plasmodesmal immune signaling cascades. Proceedings of the National Academy of Sciences. 2023b:120(17):e2216397120. 10.1073/PNAS.2216397120

64. Tee EE, Samwald S, and Faulkner C. Quantification of Cell-to-Cell Connectivity Using Particle Bombardment. 2022:263–272. 10.1007/978-1-0716-2132-5_17

65. Thomas CL, Bayer EM, Ritzenthaler C, Fernandez-Calvino L, and Maule AJ. Specific Targeting of a Plasmodesmal Protein Affecting Cell-to-Cell Communication. PLoS Biol. 2008:6(1):e7. 10.1371/JOURNAL.PBIO.0060007

66. Tomczynska I, Stumpe M, Doan TG, Mauch F. A Phytophthora effector protein promotes symplastic cell-to-cell trafficking by physical interaction with plasmodesmata-localised callose synthases. New Phytol. 2020 Sep;227(5):1467–1478. doi: 10.1111/nph.16653. Epub 2020 Jun 8. PMID: 32396661.

67. Vatén A, Dettmer J, Wu S, Stierhof YD, Miyashima S, Yadav SR, Roberts CJ, Campilho A, Bulone V, Lichtenberger R, et al. Callose Biosynthesis Regulates Symplastic Trafficking during Root Development. Dev Cell. 2011:21(6):1144– 1155. 10.1016/j.devcel.2011.10.006

68. Velásquez AC, Oney M, Huot B, Xu S, and He SY. Diverse mechanisms of resistance to Pseudomonas syringae in a thousand natural accessions of Arabidopsis thaliana. New Phytologist. 2017:214(4):1673–1687. 10.1111/NPH.14517

69. Vlot AC, Liu PP, Cameron RK, Park SW, Yang Y, Kumar D, Zhou F, Padukkavidana T, Gustafsson C, Pichersky E, et al. Identification of likely orthologs of tobacco salicylic acid-binding protein 2 and their role in systemic acquired resistance in Arabidopsis thaliana. The Plant Journal. 2008:56(3):445–456. 10.1111/J.1365-313X.2008.03618.X

70. Wang C, Dai S, Zhang ZL, Lao W, Wang R, Meng X, and Zhou X. Ethylene and salicylic acid synergistically accelerate leaf senescence in Arabidopsis. J Integr Plant Biol. 2021a:63(5):828–833. 10.1111/JIPB.13075/SUPPINFO

71. Wang X, Sager R, Cui W, Zhang C, Lu H, and Lee JY. Salicylic Acid Regulates Plasmodesmata Closure during Innate Immune Responses in Arabidopsis. Plant Cell. 2013:25(6):2315. 10.1105/TPC.113.110676

72. Wang Y, Li X, Fan B, Zhu C, and Chen Z. Regulation and Function of Defense-Related Callose Deposition in Plants. Int J Mol Sci. 2021b:22(5):1–15. 10.3390/IJMS22052393

73. Wickham H. ggplot2: Elegant Graphics for Data Analysis (Spring-Verlag New York).

74. Wickham H, Averick M, Bryan J, Chang W, D’ L, Mcgowan A, François R, Grolemund G, Hayes A, Henry L, et al. Welcome to the Tidyverse. J Open Source Softw. 2019:4(43):1686. 10.21105/JOSS.01686

75. Wildermuth MC, Dewdney J, Wu G, and Ausubel FM. Isochorismate synthase is required to synthesize salicylic acid for plant defence. Nature 2001 414:6863. 2001:414(6863):562–565. 10.1038/35107108

76. Williams LE, Lemoine R, and Sauer N. Sugar transporters in higher plants – a diversity of roles and complex regulation. Trends Plant Sci. 2000:5(7):283–290. 10.1016/S1360-1385(00)01681-2

77. Wilson DC, Kempthorne CJ, Carella P, Liscombe DK, and Cameron RK. Age-related resistance in arabidopsis thaliana involves the MADS-domain transcription factor SHORT VEGETATIVE PHASE and direct action of salicylic acid on pseudomonas syringae. Molecular Plant-Microbe Interactions. 2017:30(11):919–929. 10.1094/MPMI-07-17-0172-R/ASSET/IMAGES/LARGE/MPMI-07-17-0172-R_F8.JPEG

78. Wu S, O’Lexy R, Xu M, Sang Y, Chen X, Yu Q, Gallagher KL. Symplastic signaling instructs cell division, cell expansion, and cell polarity in the ground tissue of Arabidopsis thaliana roots. Proc Natl Acad Sci U S A. 2016 Oct 11;113(41):11621–11626. doi: 10.1073/pnas.1610358113. Epub 2016 Sep 23. PMID: 27663740; PMCID: PMC5068303.

79. Xu B, Cheval C, Laohavisit A, Hocking B, Chiasson D, Olsson TSG, Shirasu K, Faulkner C, and Gilliham M. A calmodulin-like protein regulates plasmodesmal closure during bacterial immune responses. New Phytologist. 2017:215:77–84. 10.1111/nph.14599

80. Yadav SR, Yan D, Sevilem I, Helariutta Y. Plasmodesmata-mediated intercellular signaling during plant growth and development. Front Plant Sci. 2014 Feb 17;5:44. doi: 10.3389/fpls.2014.00044. PMID: 24596574; PMCID: PMC3925825.

81. Yang L, Li B, Zheng XY, Li J, Yang M, Dong X, He G, An C, and Deng XW. Salicylic acid biosynthesis is enhanced and contributes to increased biotrophic pathogen resistance in Arabidopsis hybrids. Nature Communications 2015 6:1. 2015:6(1):1–12. 10.1038/ncomms8309

82. Zhang Z, Ruan YL, Zhou N, Wang F, Guan X, Fang L, Shang X, Guo W, Zhu S, and Zhang T. Suppressing a Putative Sterol Carrier Gene Reduces Plasmodesmal Permeability and Activates Sucrose Transporter Genes during Cotton Fiber Elongation. Plant Cell. 2017:29(8):2027–2046. 10.1105/TPC.17.00358

83. Zhongpeng L, Liu SL, Montes-Serey C, Walley JW, and Aung K. PLASMODESMATA-LOCATED PROTEIN 6 regulates plasmodesmal function in Arabidopsis vasculature. Plant Cell. 2024 koae166. 10.1093/plcell/koae166

84. Zuo J, Niu QW, and Chua NH. Technical advance: An estrogen receptor-based transactivator XVE mediates highly inducible gene expression in transgenic plants. Plant J. 2000:24(2):265–273. 10.1046/J.1365-313X.2000.00868.X

