## Supplementary Figures for "Plasmodesmal closure elicits stress responses"

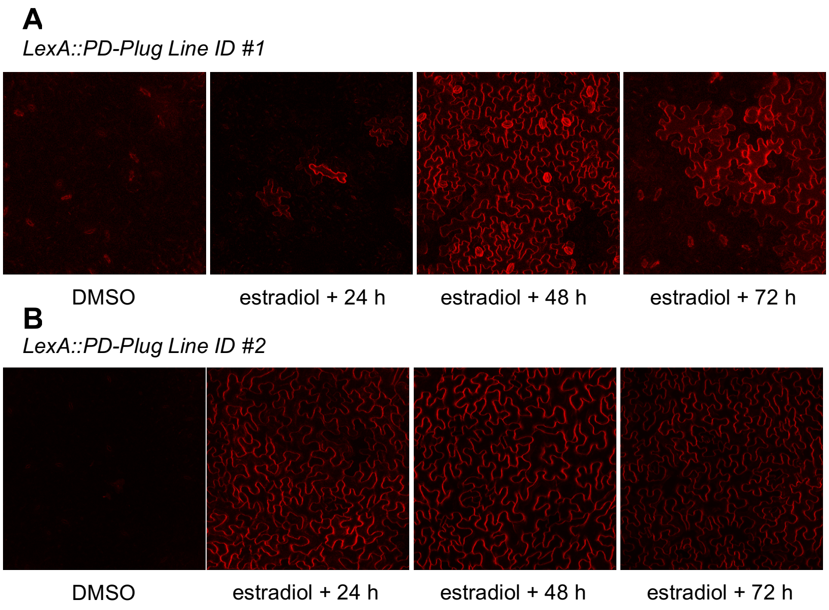


**Supplemental Figure S1. Induction of LexA::PD-Plug transgene construct after estradiol treatment in T2 independent lines.** Independent T2 lines of LexA::PD-Plug with mCherry expression showing induction post estradiol treatment. Line 2 was selected for further experimental analysis.

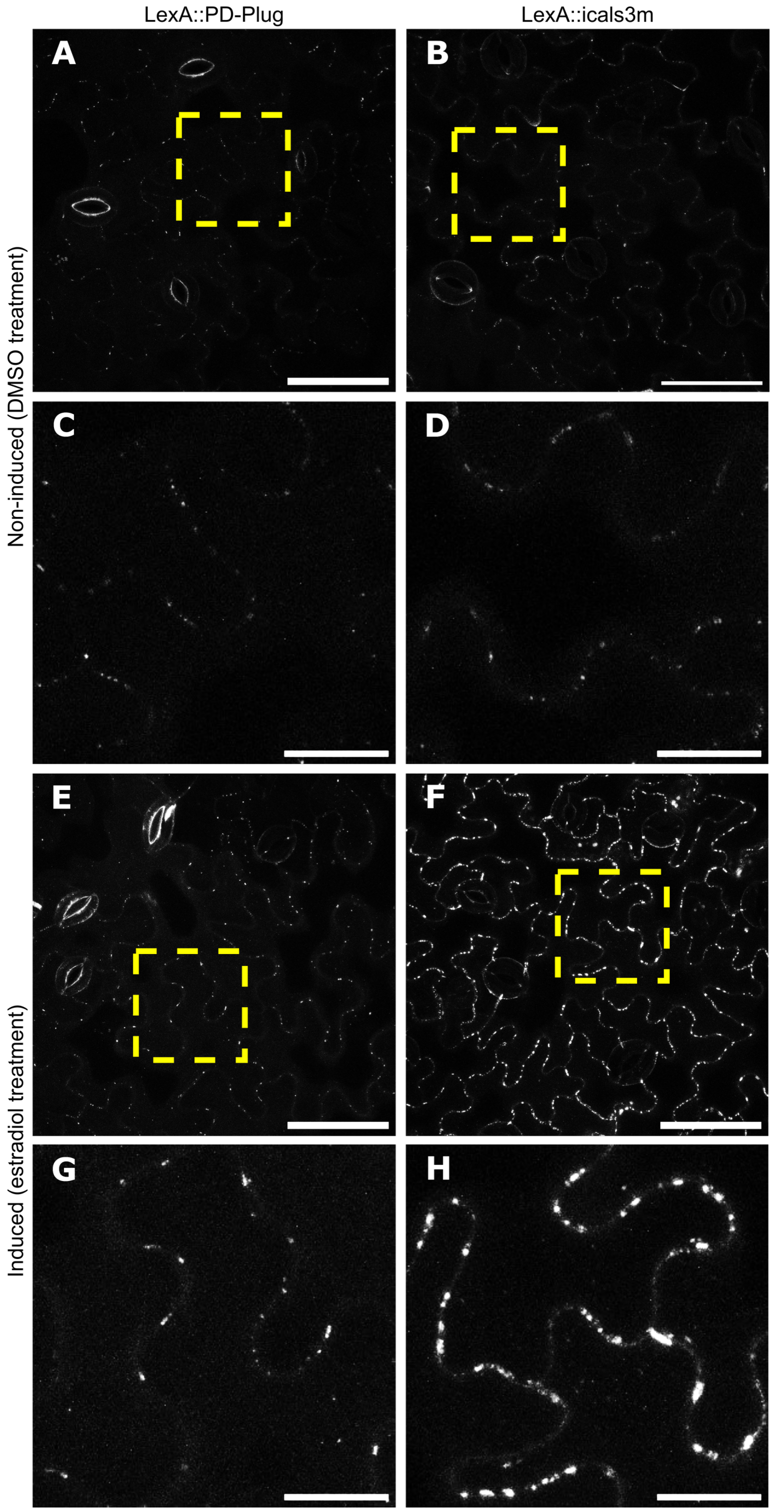


**Supplemental Figure S2. Aniline blue stained callose at plasmodesmata in LexA::PD-Plug and LexA::icals3m.** LexA::PD-Plug and LexA::icals3m treated with either DMSO (**A-D**) or estradiol (**E-H**) and infiltrated with aniline blue 24 h post treatment. **C** and **D** are zoomed in portions of **A-B**, and **G-H** is zoomed in portion of **E-F** as indicated by the yellow square. A-B and E-F scale bar = 50 μM; C-D and G-H scale bar = 15 μM.

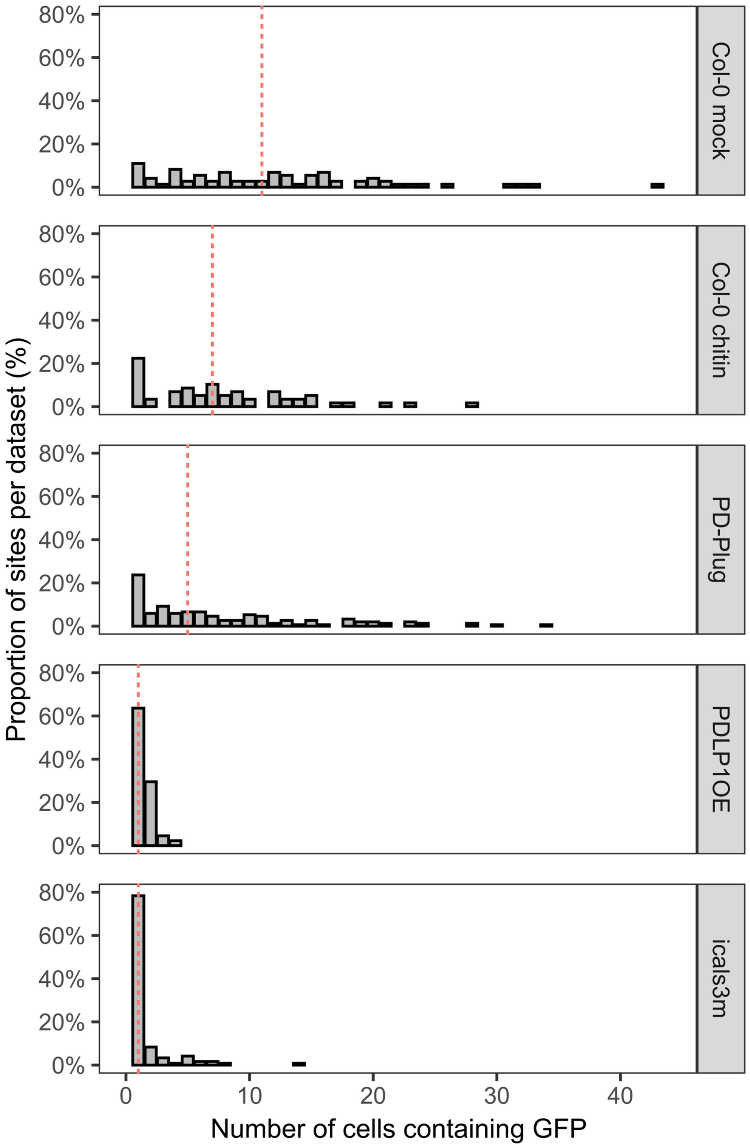


**Supplemental Figure S3. Microprojectile bombardment data in different conditions and genotypes.** Comparison of microprojectile bombardment data showing GFP movement into neighboring cells, with datasets showing high variance and heterogeneity (i.e., Col-0 mock, Col-0 chitin and estradiol treated LexA::PD-Plug [PD-Plug]) in comparison to low variance and little GFP movement (i.e., PDLP1OE and estradiol treated LexA::icals3m [icals3m]). Data represented taken from Cheval *et al*. (2020) for Col-0 mock and Col-0 chitin, and Tee *et al*. (2023) for PDLP1OE. Red dotted line indicates median in a given dataset.

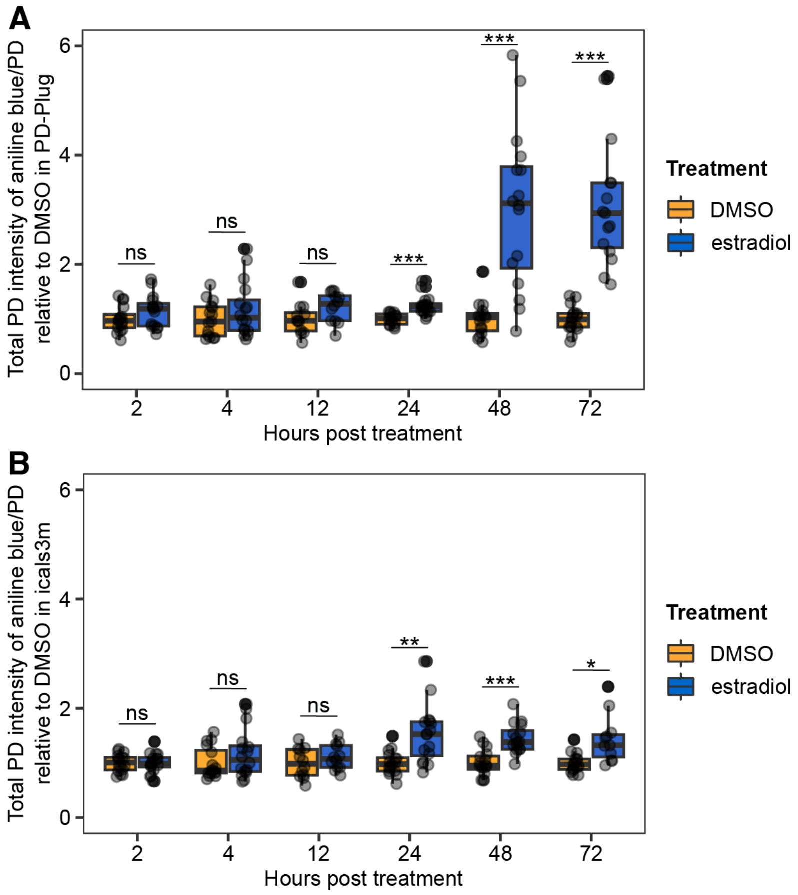


**Supplemental Figure S4. Time course callose quantification in LexA::PD-Plug and LexA::icals3m treated with DMSO or estradiol.** Aniline blue-stained callose deposition at plasmodesmata quantification in either LexA::PD-Plug (**A**) or LexA::icals3m (**B**). Bootstrap analysis indicates significant differences between DMSO and estradiol treatment at each time point, as indicated by * $p$<0.05, ** $p$<0.01, or *** $p$<0.001, with n ≥ 11 images with a minimum of 3 biological replicates per genotype/treatment/timepoint.

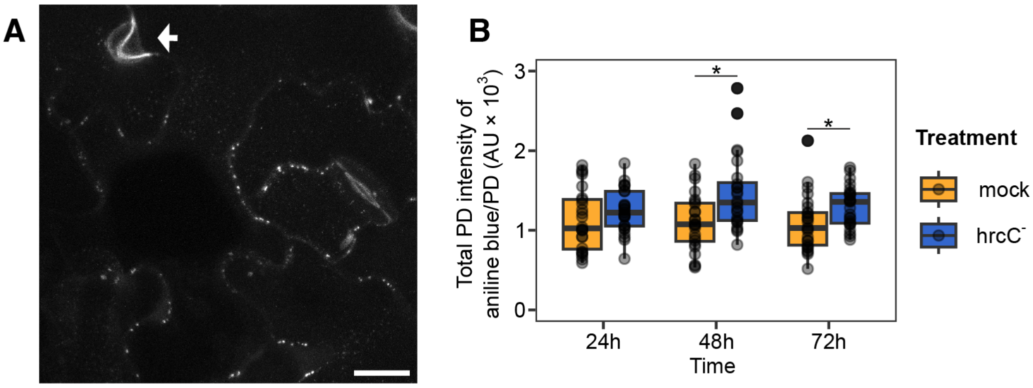


**Supplemental Figure S5. *Pseudomonas syringae* DC3000 mutant strain hrcC$^{-}$ infection in Col-0. A)** Example aniline blue-stained callose deposition at plasmodesmata as well as macroscopic callose deposits (labelled by white arrow) with the *Pseudomonas syringae hrcC* mutant (*hrcC*) treatment. Scale bar = 15 µm. **B)** Quantification of aniline blue-stained plasmodesmata-associated callose in 5-week-old Col-0 plants 24 h, 48 h and 72 h post treatment of $H_2O$ (mock) or *hrcC*. Datapoints represent the average of the total PD intensity/plasmodesmata (PD) per image, with n ≥ 24 images from 8 biological replicates per treatment/timepoint. Bootstrap analysis indicates significant differences between mock and *hrcC* treatment at 48 h and 72 h, as indicated by * $p<0.05$.

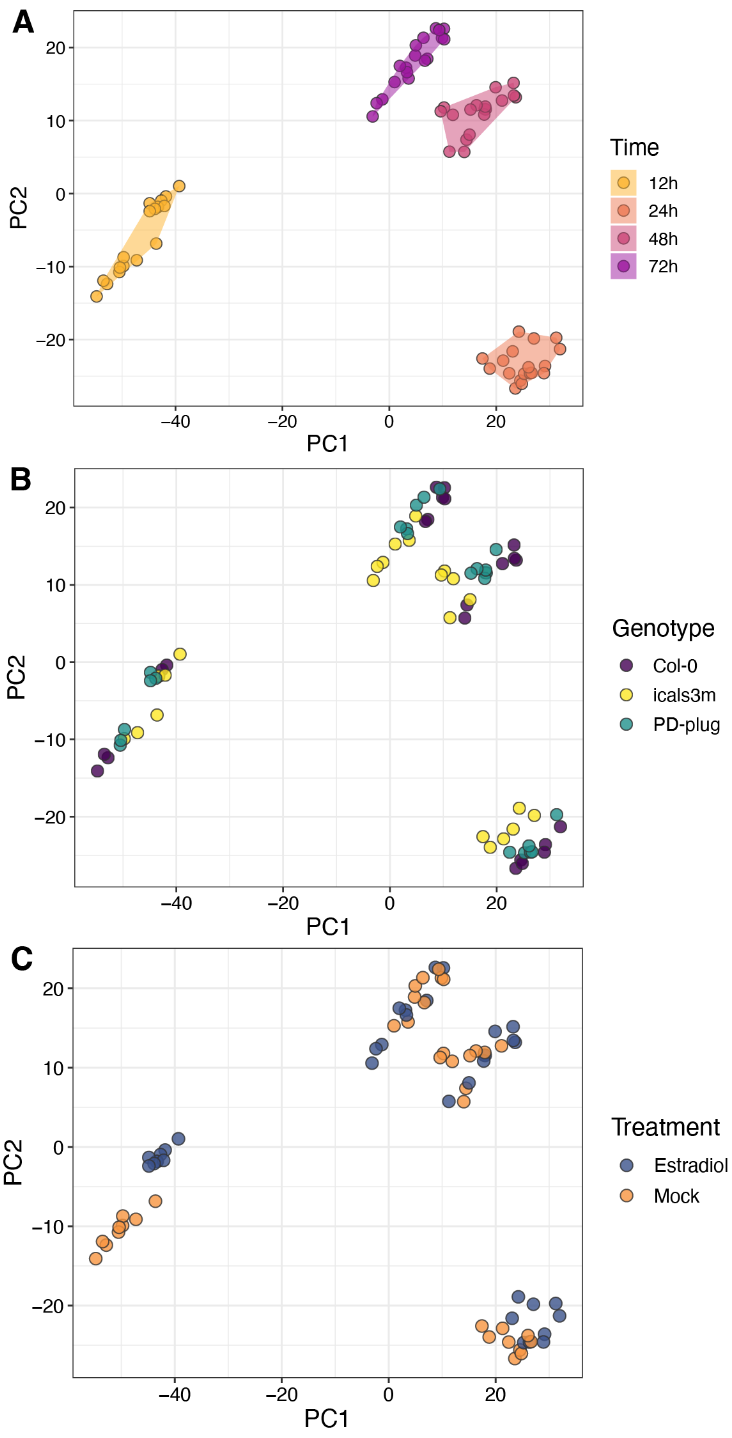


**Supplemental Figure S6. PCA of genotypes Col-0, LexA::PD-Plug and LexA::icals3m treated with DMSO or estradiol at each time point.** PCA grouped based on time (**A**), genotype (**B**) and treatment (**C**) indicate a critical driving factor of variance was time.

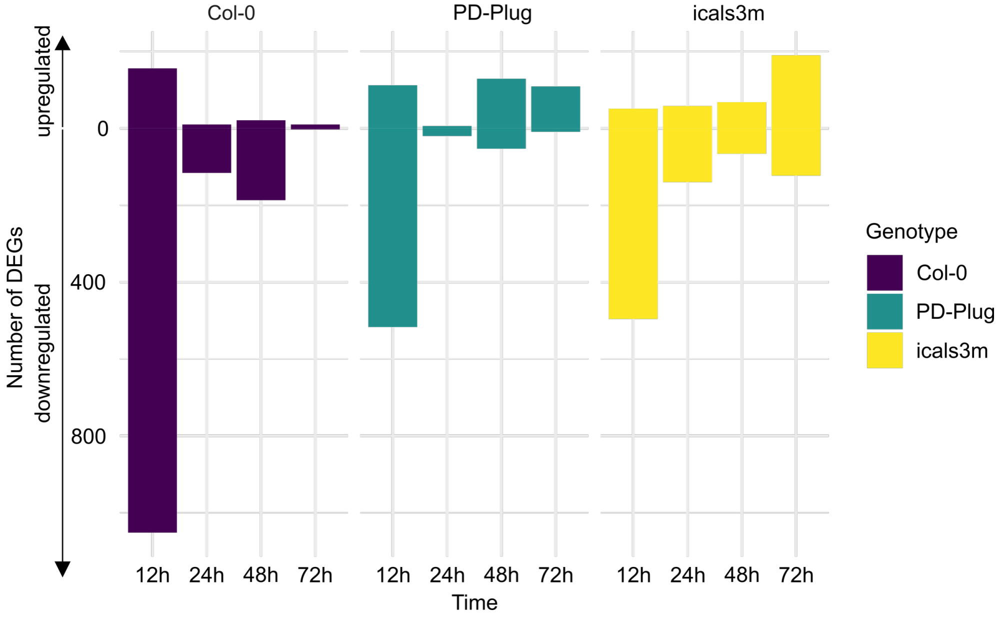


**Supplemental Figure S7. Estradiol induces transcriptional changes.** Number of differentially expressed genes (DEGs) in each genotype, when treated with estradiol compared to DMSO. 12 h after treatment, hundreds of genes are downregulated in all genotypes with a large effect seen in Col-0. The effect of estradiol in Col-0 is largely negated by 24 h (see Supplemental Data Set 1 for shared genes across genotypes), with no effect of estradiol detected in Col-0 by 72 h.

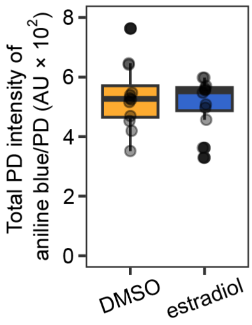


**Supplemental Figure S8. Estradiol has no effect on plasmodesmal callose accumulation Col-0 at 12 h.** Aniline blue-stained callose deposition at plasmodesmata quantification in Col-0 with either DMSO or estradiol treatment. Bootstrap analysis indicates no significant difference between treatments, with n = 12 images and 3 biological replicates per treatment.

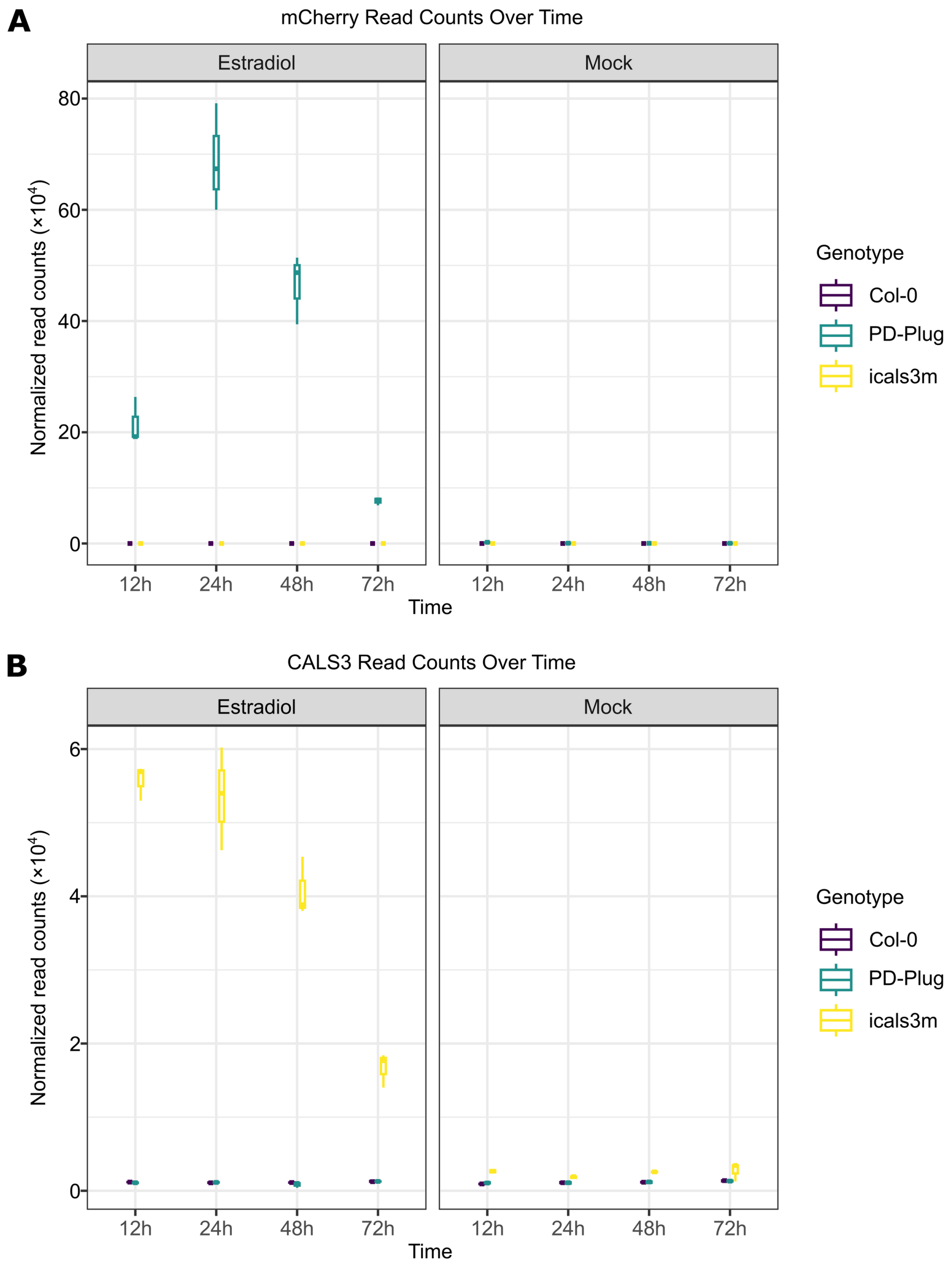


**Supplemental Figure S9. Estradiol-induced transgene expression over 72 h.** Normalized read counts of estradiol induced transgene expression at each time point for each genotype (Estradiol, left panels), in comparison to the respective DMSO treated (Mock, right panels) samples. **A)** mCherry read counts, indicative of the LexA::PD-Plug transgene; **B)** CalS3 read counts, including reads from both the native CalS3 expression and that induced by the LexA::icals3m transgene.

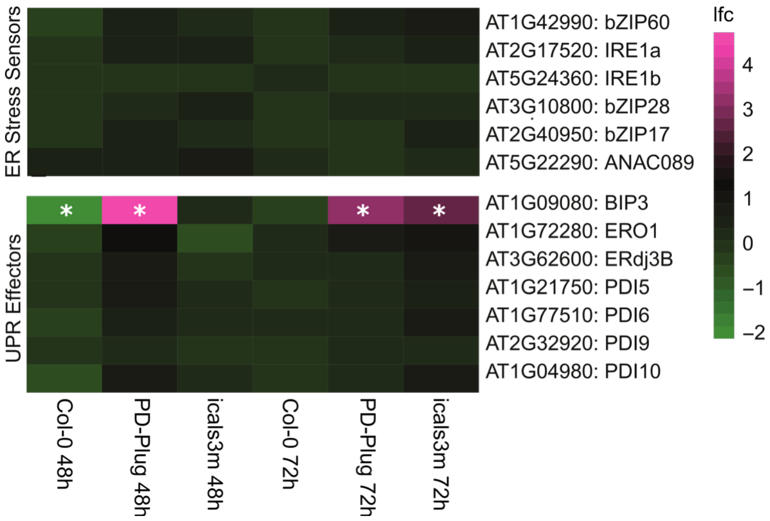


**Supplemental Figure S10. Gene expression of genes related to ER stress.** Effect of estradiol on genes defined as ER stress sensors or unfold protein response (UPR) effectors (Beaugelin et al. 2020), in genotypes Col-0, LexA::PD-Plug and LexA::icals3m. Stars indicate significant up or down-regulation in comparison to DMSO treatment.

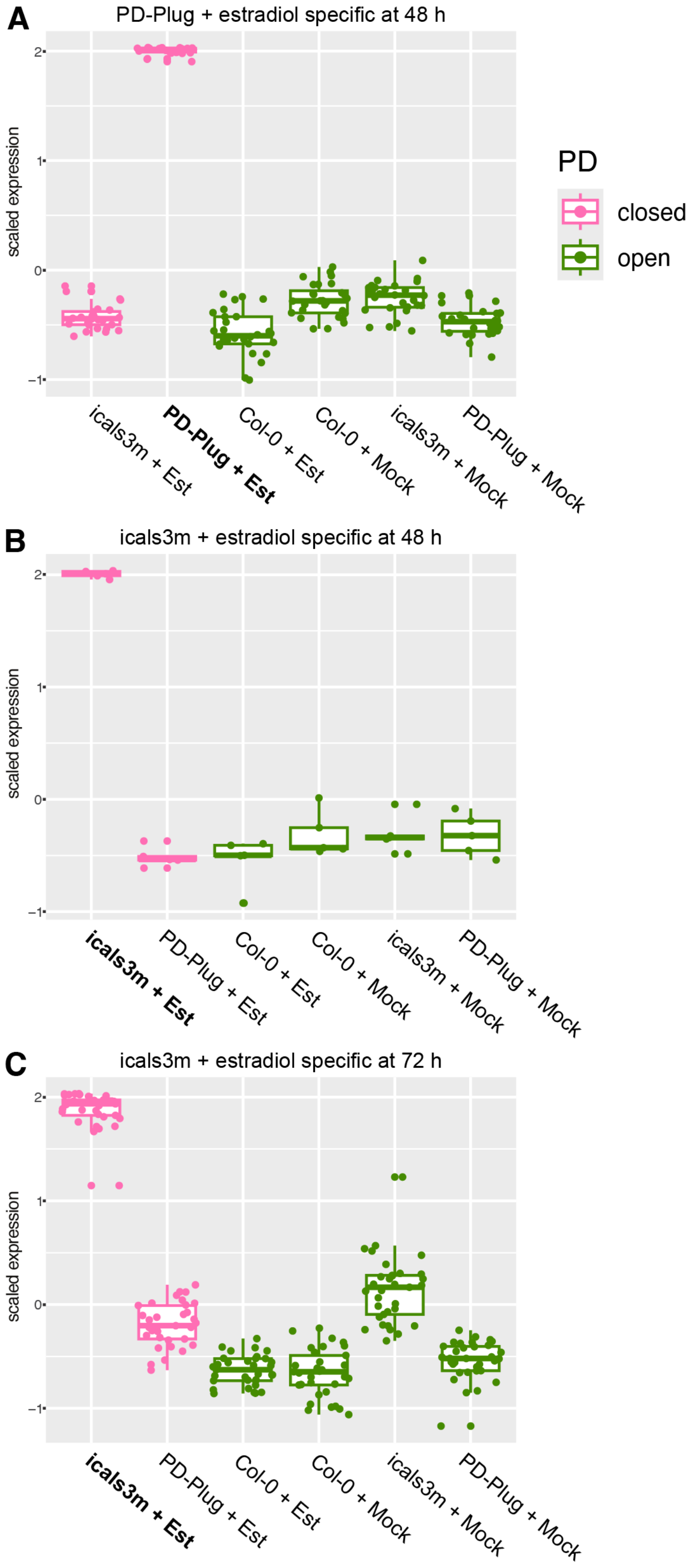


**Supplemental Figure S11. Likelihood ratio clustering for genes significantly different in estradiol treated LexA::PD-Plug at 48 h (A), estradiol treated LexA::icals3m at 48 h (B) and estradiol treated LexA::icals3m at 72 h (C).** Plasmodesmal state (PD) indicated by color, with 'closed' plasmodesmata being LexA::PD-Plug (PD-Plug) and LexA::icals3m (icals3m) treated with estradiol (Est), and ‘open’ plasmodesmata being LexA::PD-Plug and LexA::icals3m treated with DMSO (Mock), and Col-0 treated with DMSO (Mock) and estradiol (Est).

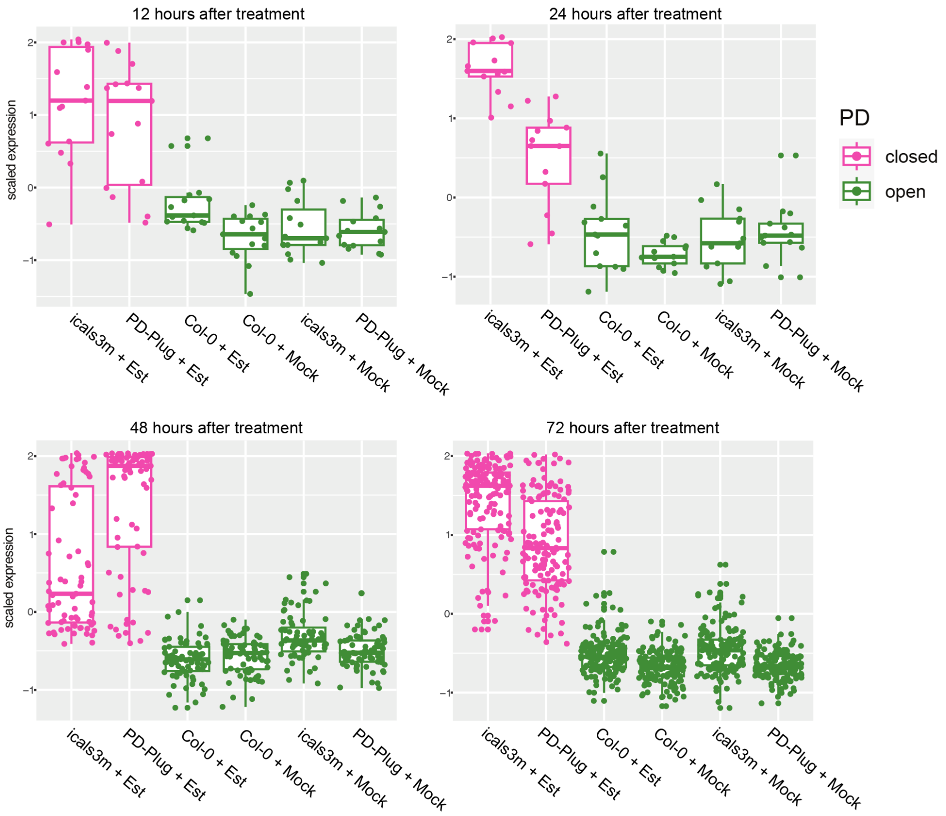


**Supplemental Figure S12. Likelihood ratio clustering of groups based on plasmodesmal state.** Groups based on plasmodesmal state (PD), with ‘closed’ plasmodesmata being LexA::PD-Plug (PD-Plug) and LexA::icals3m (icals3m) treated with estradiol (Est), and ‘open’ plasmodesmata being LexA::PD-Plug and LexA::icals3m treated with DMSO (Mock), and Col-0 treated with DMSO (Mock) and estradiol (Est).

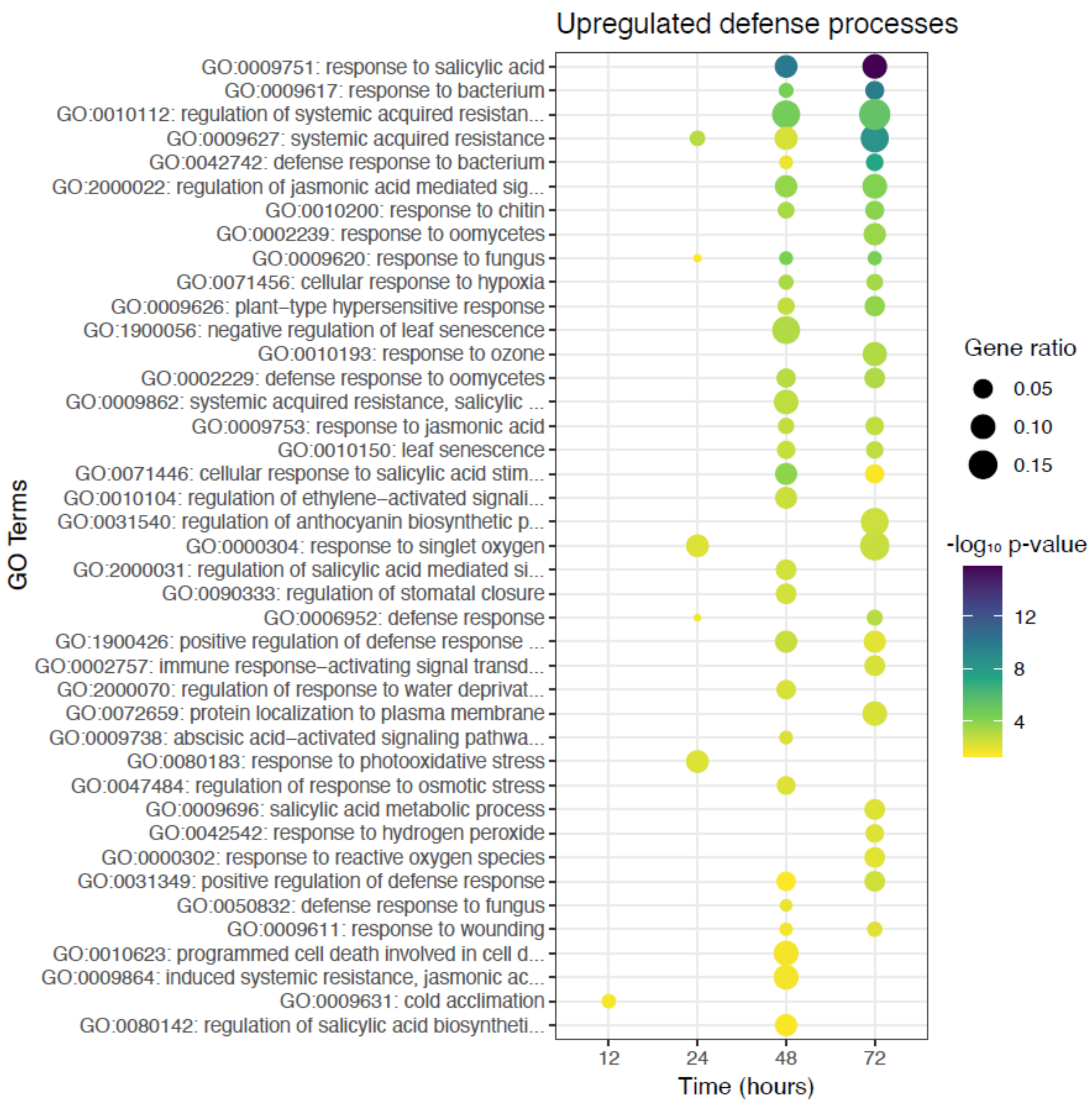


**Supplemental Figure S13. GO terms enriched when plasmodesmata are closed, under the parent term of defense processes shown across all time points.**

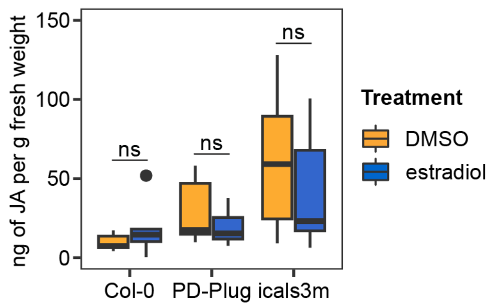


**Supplemental Figure S14. Jasmonic acid quantification in Col-0, PD-Plug, and icals3m.** Quantification of jasmonic acid in 5-week-old plants after 72 h DMSO or estradiol treatment, n = 6 per genotype/treatment. No significant differences (ns) between treatment within a genotype was found using the Mann-Whitney test.

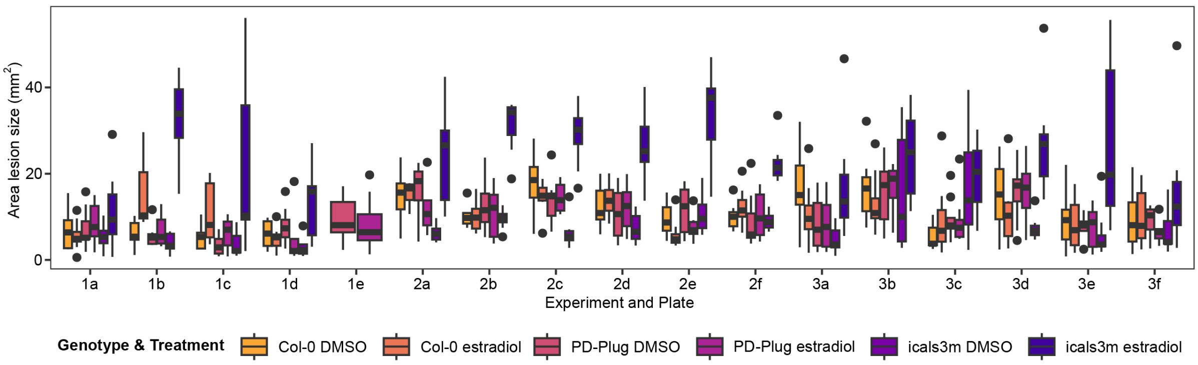


**Supplemental Figure S15. *Botrytis cinerea* disease lesions in Col-0, LexA::PD-Plug and LexA::icals3m.** Area of disease lesions 2 dpi in leaves of 5-week-old plants, inoculated after 72 h of DMSO or estradiol treatment. Graph of each plate within an experiment as indicated by the number (experiment) and letter (plate). Four leaves were represented for each treatment/genotype combination per plate.

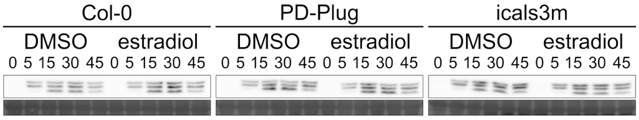


**Supplemental Figure S16. MAPK-activation by flg22 in seedlings of Col-0, LexA::PD-Plug and LexA::icals3m, 72 h post DMSO or estradiol treatment.** Western blot with minute timings denoted, and Coomassie blue staining indicated below as loading controls.

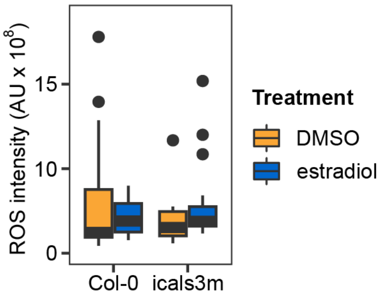


**Supplemental Figure S17. Basal ROS quantification of LexA::icals3m in comparison to Col-0.** Quantification of ROS from z-stacks using sum projection, as indicated utilizing 20 µM $H_2$DCFDA. 5-week-old plants were treated with DMSO or estradiol for 72 h before visualization, with data from four images per plant, n = 4 per treatment/genotype. Independent factors genotype and treatment were not significant, with no significant interaction between the two (ANOVA; F = 0.05, df = 2, P = 0.83; F = 0.002, df = 1, P = 0.97; F = 1.08, df = 1, P = 0.32).

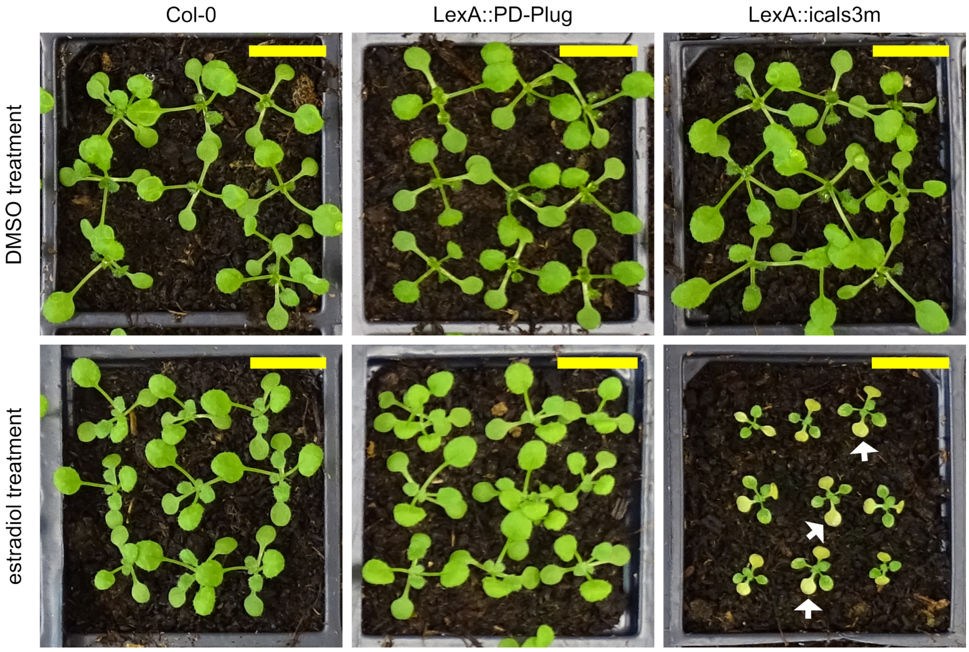


**Supplemental Figure S18. Photographs of 15-day old plants showing the growth phenotype of Col-0, LexA::PD-Plug, LexA::icals3m induced by DMSO or estradiol treatment.** White arrows indicate yellowing/senescence on leaves. Scale bar = 15 mm.

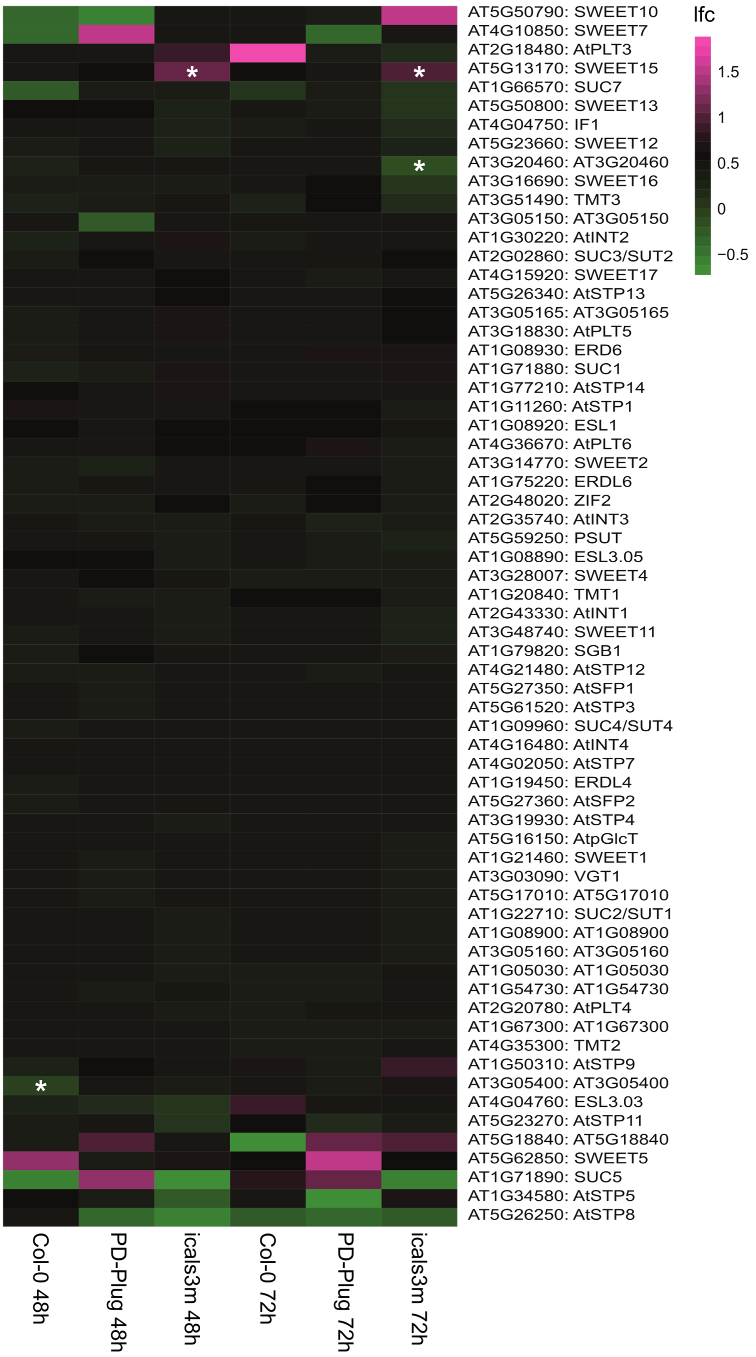


**Supplemental Figure S19. Heat map showing gene expression of sugar related genes at 48 h and 72 h post treatment.** Effect of estradiol on genes related to sugar transporters, in genotypes Col-0, LexA::PD-Plug and LexA::icals3m, with stars indicating significant up or down-regulation in comparison to DMSO treatment.

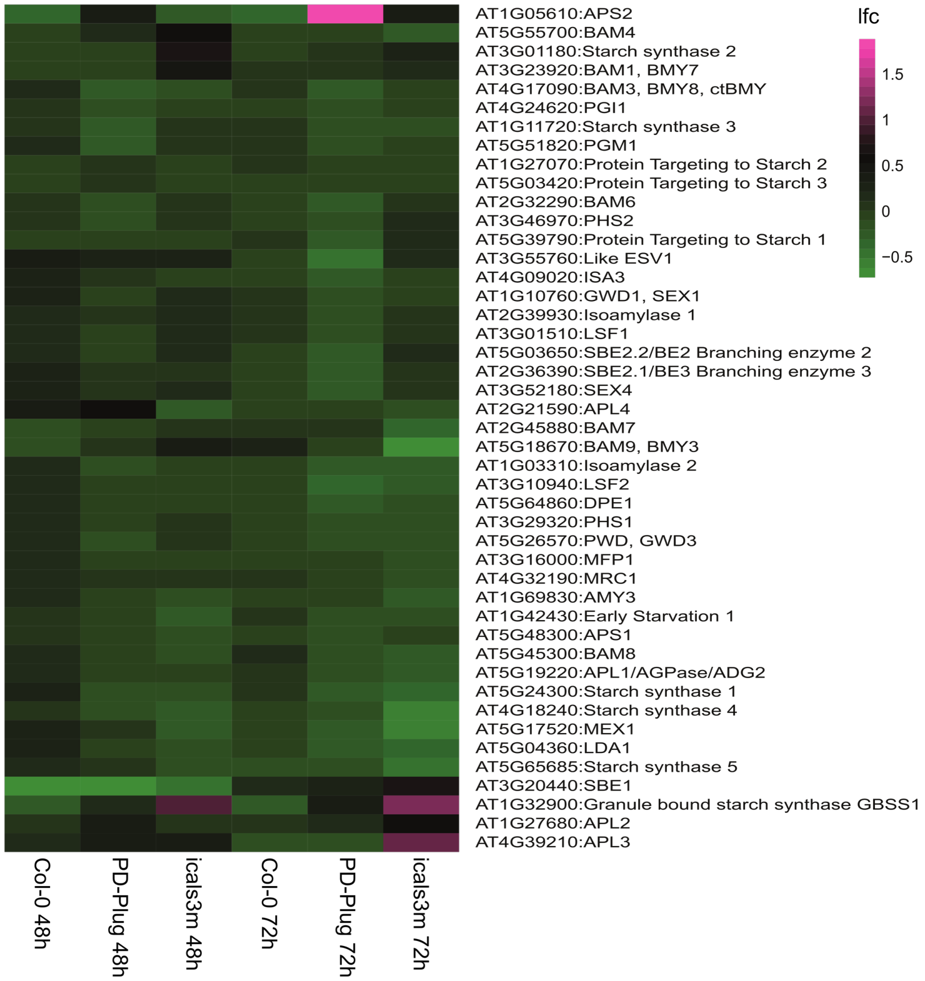


**Supplemental Figure S20. Gene expression of starch related genes at 48 h and 72 h post treatment.** Effect of estradiol on expression of genes related to starch in genotypes Col-0, LexA::PD-Plug and LexA::icals3m.

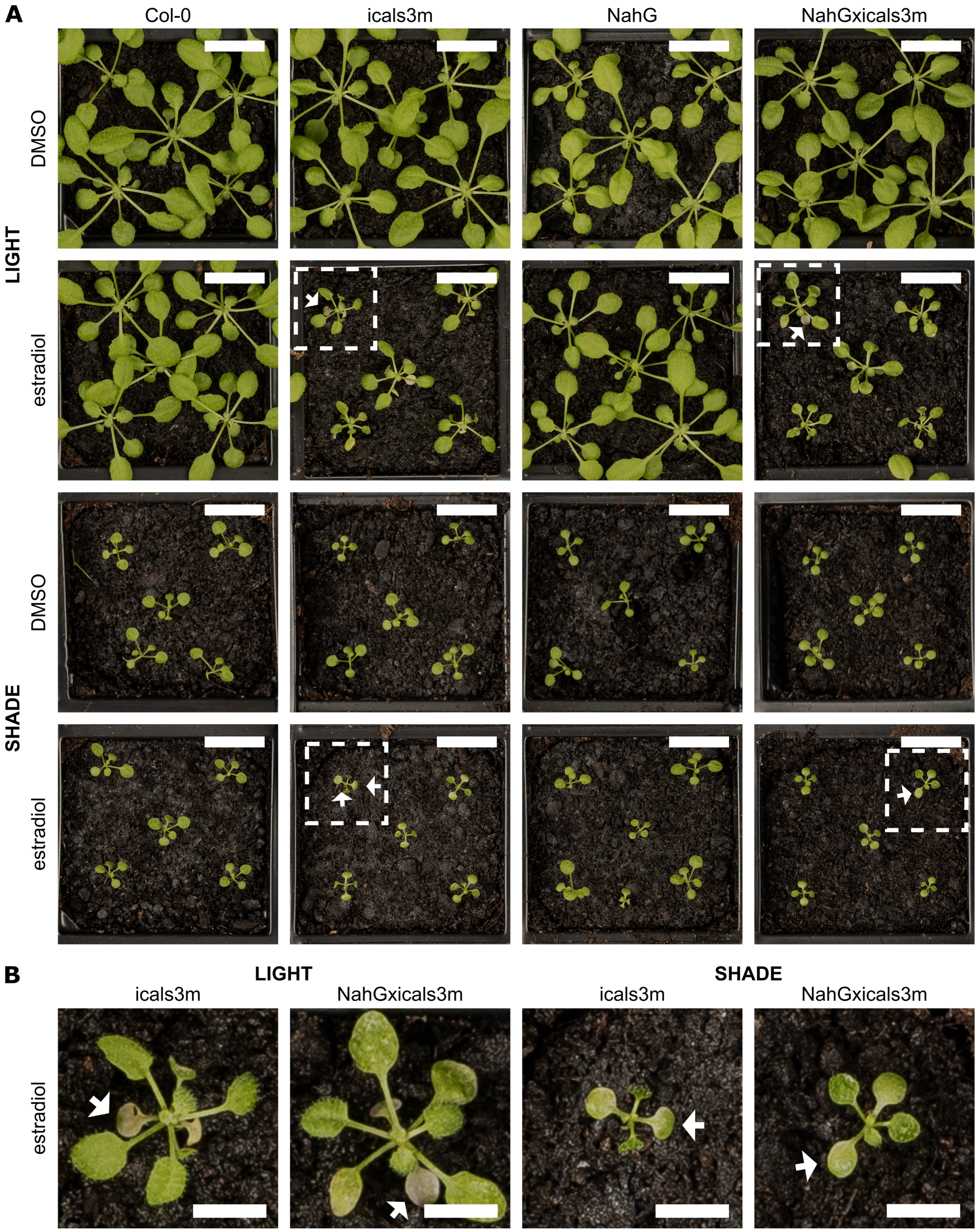


**Supplemental Figure S21. Photographs of 24-day old plants showing the growth phenotype of Col-0, LexA::icals3m (icals3m), NahG and NahG×LexA::icals3m (NahG×icals3m) induced by DMSO or estradiol treatment under light or shaded conditions.** White arrows indicate example yellowing/senescence on leaves. **B** is zoomed in portion of **A** as indicated by the white square. **A** scale bar = 15 mm, **B** scale bar = 5mm.

**Supplemental Table 1. Selection information for LexA::icals3m transgenic line.** Independent T2 lines as indicated by Line ID, were analyzed for copy number and a qPCR was performed on seedings 24 h post treatment of DMSO or estradiol, with gene expression indicated by relative normalized quantity (NRQ). Fold change difference is estradiol NRQ/DMSO NRQ. Chosen line for characterization indicated by bold and italicized text.

| Line ID | Copy Number | DMSO NRQ | Estradiol NRQ | Fold Change |
|---|---|---|---|---|
| ***#23*** | ***2*** | ***0.03*** | ***0.44*** | ***15.14*** |
| #10 | 2 | 0.02 | 0.10 | 6.05 |
| #22 | 2 | 0.02 | 0.05 | 3.19 |
| #7 | 3 | 0.03 | 0.33 | 10.73 |
| #19 | 3 | 0.07 | 0.15 | 2.09 |
| #2 | 4 | 0.02 | 0.31 | 19.52 |
| #17 | 4 | 0.06 | 0.80 | 13.93 |
| #6 | 16 | 0.07 | 2.45 | 34.94 |
| #12 | 24 | 0.16 | 1.19 | 7.26 |
